# Active transcription and Orc1 drive chromatin association of the AAA+ ATPase Pch2 during meiotic G2/prophase

**DOI:** 10.1101/777003

**Authors:** Richard Cardoso da Silva, María Ascensión Villar-Fernández, Gerben Vader

## Abstract

Pch2 is an AAA+ protein that controls DNA break formation, recombination and checkpoint signaling during meiotic G2/prophase. Chromosomal association of Pch2 is linked to these processes, and several factors influence the association of Pch2 to euchromatin and the specialized chromatin of the ribosomal (r)DNA array of budding yeast. Here, we describe a comprehensive mapping of Pch2 localization across the budding yeast genome during meiotic G2/prophase. Within non-rDNA chromatin, Pch2 associates with a subset of actively RNA Polymerase II (RNAPII)-dependent transcribed genes. Chromatin immunoprecipitation (ChIP)- and microscopy-based analysis reveals that active transcription is required for chromosomal recruitment of Pch2. Similar to what was previously established for association of Pch2 with rDNA chromatin, we find that Orc1, a component of the Origin Recognition Complex (ORC), is required for the association of Pch2 to these euchromatic, transcribed regions, revealing a broad connection between chromosomal association of Pch2 and Orc1/ORC function. Ectopic mitotic expression is insufficient to drive recruitment of Pch2, despite the presence of active transcription and Orc1/ORC in mitotic cells. This suggests meiosis-specific ‘licensing’ of Pch2 recruitment to sites of transcription, and accordingly, we find that the synaptonemal complex (SC) component Zip1 is required for the recruitment of Pch2 to transcription-associated binding regions. Interestingly, Pch2 binding patterns are distinct from meiotic axis enrichment sites (as defined by Red1, Hop1 and Rec8). This suggests that although Pch2 is linked to axis/SC-directed recruitment and function, the chromosomal population of Pch2 described here is not directly associated with chromosomal axis sites. In line with this observation, interfering with the pool of Pch2 that associates with active RNAPII transcription does not lead to effects on the chromosomal abundance of Hop1, a known axial client of Pch2. We thus report characteristics and dependencies for Pch2 recruitment to meiotic chromosomes, and reveal an unexpected link between Pch2, SC formation, chromatin and active transcription.

## Introduction

Meiosis is a specialized developmental program dedicated to the production of genetically unique haploid gametes [1]. The production of haploid gametes is made possible by several meiosis-specific events, chief among them the event of homologous chromosome segregation during the first meiotic chromosome segregation event (*i.e.* meiosis I). Faithful segregation of homologs requires that initially unconnected homologous chromosomes are physically linked prior to segregation. Homolog linkage is achieved by interhomologue-directed crossover repair of programmed DNA double strand breaks (DSBs) prior to meiosis I (*i.e.* during meiotic G2/prophase). DSBs are introduced by Spo11, a topoisomerase-like protein, which acts in conjunction with several accessory factors [2]. DSB formation happens in the context of a specialized, meiosis-specific chromosome architecture [3] [4]. Several protein factors (such as Red1 and Hop1 in budding yeast [5] [6]) drive the assembly of chromosomes into linear arrays of chromatin loops that emanate from a proteinaceous structure termed the meiotic chromosome axis. Red1 and Hop1 co-localize with the meiotic cohesin complex (containing the meiosis-specific Rec8 kleisin subunit instead of the canonical Scc1) to form the molecular foundation of this typical meiotic ‘axis-loop’ chromosome structure [7, 8]. A zipper-like assembly called the synaptonemal complex (SC) polymerizes between synapsing homologous chromosomes [9], concomitantly with, and dependent on ongoing crossover repair of meiotic DSBs [10, 11]. In budding yeast, the Zip1 protein is the main component of the SC, which is assembled onto the axial components of the loop-axis architecture [12, 13]. The SC likely acts as a signaling conduit that coordinates DSB activity and repair template preferences with chromosome synapsis [14-16]. A major role for the SC lies in directing the chromosomal recruitment of the hexameric AAA+ enzyme Pch2 [14, 17, 18], an important mediator of DSB activity, repair, and checkpoint function (reviewed in [19]). The molecular mechanisms of Pch2 recruitment to synapsed chromosomes remain poorly understood. In *zip1Δ* cells, Pch2 cannot be recruited to meiotic chromosomes (except to the nucleolus/rDNA; see below) [17]. However, this is unlikely via a direct molecular interaction. First, a specific Zip1-mutant (*zip1-4LA*) uncouples SC formation from Pch2 recruitment [14, 20]. Second, in cells lacking the histone H3 methyltransferase Dot1, Pch2 can be recruited to unsynapsed chromosomes in *zip1Δ* cells [21, 22]. Third, a recent report has linked topoisomerase II (Top2) function to Pch2 association with synapsed chromosomes [23], hinting at a connection between chromosome topology and Pch2 recruitment.

Functionally, a main role for Pch2 on synapsed chromosome has been identified in modulating the abundance of Hop1 on chromosomes [14, 17, 24]. Pch2 recruitment to SC-forming chromosomal regions allows it to use its ATPase activity to dislodge Hop1 from synapsed regions [25-27], causing a coupling of SC formation to a reduction in DSB activity, interhomologue repair bias and checkpoint function [14, 19]. In addition to its recruitment to euchromatic regions, Pch2 is recruited to the nucleolus, where it is involved in protecting specific regions of the ribosomal (r)DNA array (and rDNA-flanking euchromatic regions) against Spo11-directed DSB activity [17, 28]. The nucleolus is devoid of SC polymerization (and thus of Zip1), and nucleolar recruitment of Pch2 is dependent on Sir2 (a histone deacetylase) and Orc1 (a component of the Origin Recognition Complex (ORC)) [17, 28]. Strikingly, with the exception of Zip1, all factors that direct Pch2 recruitment (whether within the rDNA, or within euchromatin) are involved in chromatin function, be it modification (Dot1 and Sir2), binding (Orc1, via its bromo-adjacent homology (BAH) domain) or metabolism (Topoisomerase II). Together, these observations predict an intimate interplay between chromatin and Pch2 binding. Inspired by this, and with the aim of increasing our understanding of Pch2 function on meiotic chromosomes, we generated a comprehensive map of Pch2 chromosomal association during meiotic G2/prophase. This analysis revealed specific binding sites of Pch2 across the genome. Within euchromatin, these sites map to regions of RNA Polymerase II (RNAPII)-driven transcriptional activity (*i.e.* a subset of active genes), and recruitment of Pch2 depended on active RNAPII-driven transcription. Orc1 (and also other ORC subunits) are enriched at Pch2 binding sites, whereas no Pch2 can be found associated with origins of replication, which are the canonical binding sites of ORC [29]. Intriguingly, Orc1 inactivation triggers loss of Pch2 binding at active genes, demonstrating a connection between Pch2 and Orc1 that extends beyond their previously described shared rDNA-associated functions [28]. Although active transcription and Orc1 are equally present in meiotic and mitotic cells, we further show that ectopic expression of Pch2 in vegetatively growing cells is not sufficient to allow recruitment of Pch2 to the identified binding sites within actively transcribed genes. This suggests meiosis-specific requirements that license Pch2 recruitment. In agreement with this, we find that Zip1 is required for the recruitment of Pch2 to the identified transcription-associated binding regions.

Interestingly, the Pch2 binding patterns identified here are distinct from meiotic axis enrichment sites (as defined by Red1, Hop1 and Rec8). Thus, although Pch2 has been associated with axis/SC-directed recruitment and function, the chromosomal population of Pch2 identified here is likely not directly associated with chromosomal axis sites. In line with this observation, we find that interfering with the pool of Pch2 that associates with active RNAPII transcription does not lead to effects on the chromosomal association of Hop1. We thus uncover characteristics and dependencies for Pch2 recruitment to meiotic chromosomes, and reveal an unexpected link between Pch2, SC formation, chromatin and active transcription.

## Material and Methods

### Yeast strains and growth conditions

All yeast strain used in this study were of the SK1 strain background, except for the strains harboring the galactose-inducible promoter (*pGAL10*) system which are of the W303 background. The genotypes of these strains are listed in Supplementary Data. Induction of synchronous meiosis was performed as described in [28]. For experiments using mitotically cycling cells (as shown in Figure 5A-D), cells were grown to saturation in YP-D/R medium ((1% (w/v) yeast extract, 2%(w/v) peptone, 0,1%(w/v) dextrose and 2%(w/v) raffinose)) at 30°C. Cultures were diluted to an optical density at 600nm (OD600) of 0.4, grown for an additional 4 hours after which 2% galactose was added. Unless stated otherwise, samples of cells undergoing synchronous meiosis were collected 4 hours after incubation in sporulation (SPO) medium. Synchronous entry of cultures into the meiotic program was confirmed by flow cytometry-based DNA content analysis (see below). For experiments using temperature sensitive strains, meiotic induction was performed as described in [28], except that cells were grown for up to 24 hours in pre-sporulation medium (BYTA) at the permissive temperature (23°C). Meiotic cultures were kept at 23°C (for *orc1-161*strains) or shifted to 30°C (for *orc2-1* strains). For the inhibition of global transcription (Supplementary Figure 3) 1,10-Phenanthroline (100 μg/mL in 20% ethanol, Sigma-Aldrich) [30] was added to cultures 3 hours after induction into the meiotic program. Cells were subsequently grown for one hour, and harvested. For mitotic expression of Pch2-E399Q, the coding sequence of *pch2*-E399Q (lacking its intron) was cloned in a *URA3* integrative plasmid containing *pGAL10-3XHA.* The plasmid was integrated at the *URA3* locus. For expression of 3XFLAG-dCas9 in meiosis, *3XFLAG-dCas9-tCYC1* was cloned in a TRP1 integrative plasmid containing *pHOP1* to create *pHOP1*-*3XFLAG-dCas9-tCYC1.* The plasmid containing *3XFLAG-dCas9/pTEF1p-tCYC1* was a gift from Hodaka Fujii and obtained via Addgene.org (Addgene plasmid #62190) [31].

**Figure 1.**
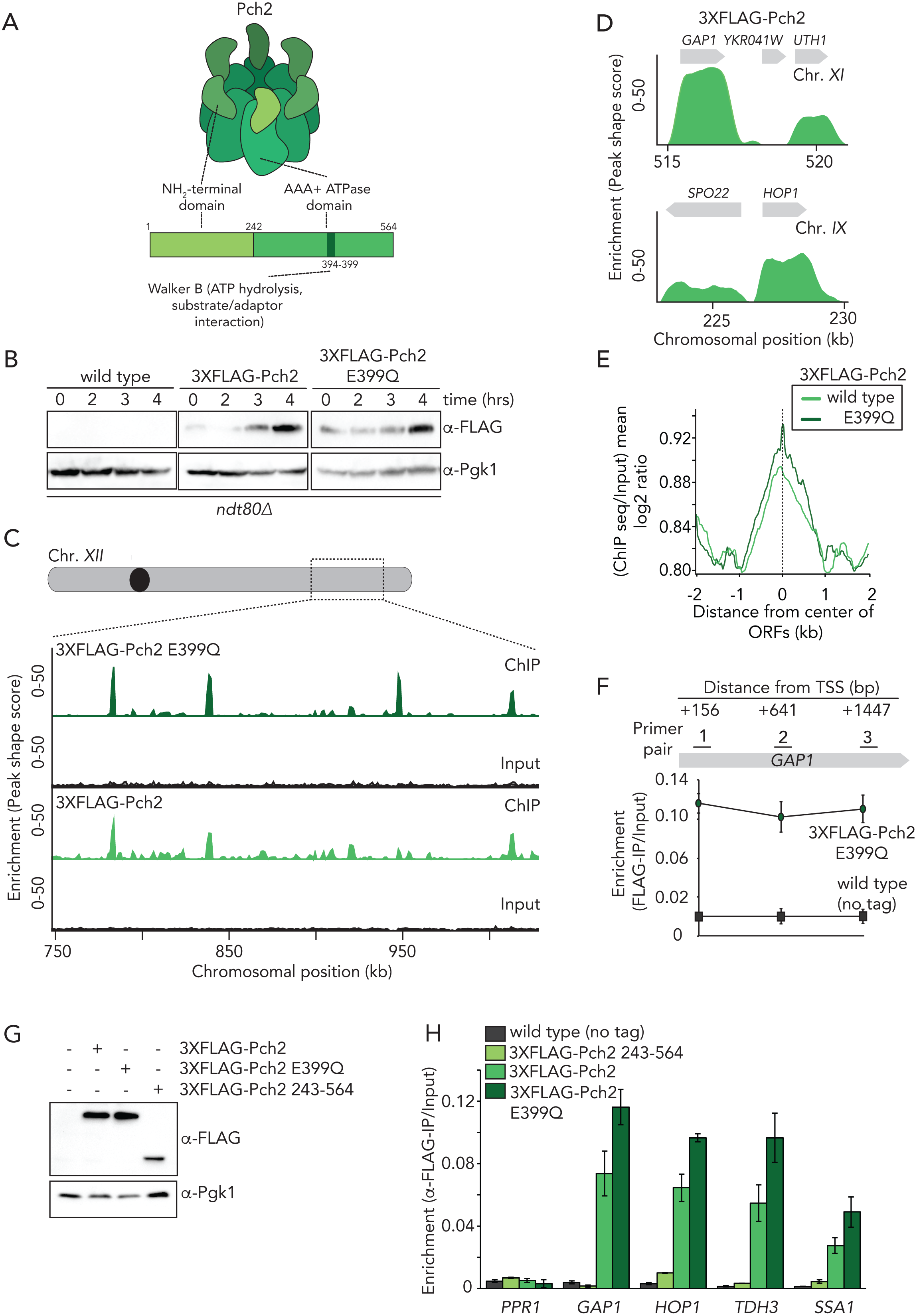
Genome-wide analysis of Pch2 chromosome association. A. Schematic of Pch2 domain organization. B. Western blot analysis of expression of 3XFLAG-Pch2 and 3XFLAG-Pch2-E399Q during meiotic G2/prophase. Time (hours) after induction into the meiotic program are indicated. C. Representative image of ChIP-seq binding patterns (ChIP and input) for 3XFLAG-Pch2 and 3XFLAG-Pch2-E399Q. Shown is a region of Chromosome *XII* (chromosomal coordinates (kb) are indicated). D. High resolution examples of 3XFLAG-Pch2 binding patterns across two selected chromosomal regions. Chromosomal coordinates and gene organization are indicated. E. Whole genome average plotting of 3XFLAG-Pch2 and 3XFLAG-Pch2-E399Q binding peaks (log2). Datasets were aligned relative to the center of ORFs. F. ChIP-qPCR analysis of three locations along the *GAP1* locus during meiotic G2/prophase (4 hours). Primers pairs 1: yGV2595/yGV2596, 2: yGV2597/yGV2598, 3: yGV2599/yGV2600. Error bars represent standard error of at least three independent experiments performed in triplicate. G. Western blot analysis of expression of 3XFLAG-Pch2, 3XFLAG-Pch2-E399Q and 3XFLAG-*Δ*NTD-Pch2 during meiotic G2/prophase (4 hours). H. ChIP-qPCR analysis of 3XFLAG-Pch2, 3XFLAG-Pch2-E399Q and 3XFLAG-*Δ*NTD-Pch2 at *PPR1* (yGV2390/yGV2391), *GAP1* (yGV2597/yGV2598), *HOP1*(yGV2607/yGV2608), *TDH3* (yGV2591/yGV2592), and *SSA1* (yGV2587/yGV2588) during meiotic G2/prophase (4 hours). Error bars represent standard error of at least three independent experiments performed in triplicate.

**Figure 2.**
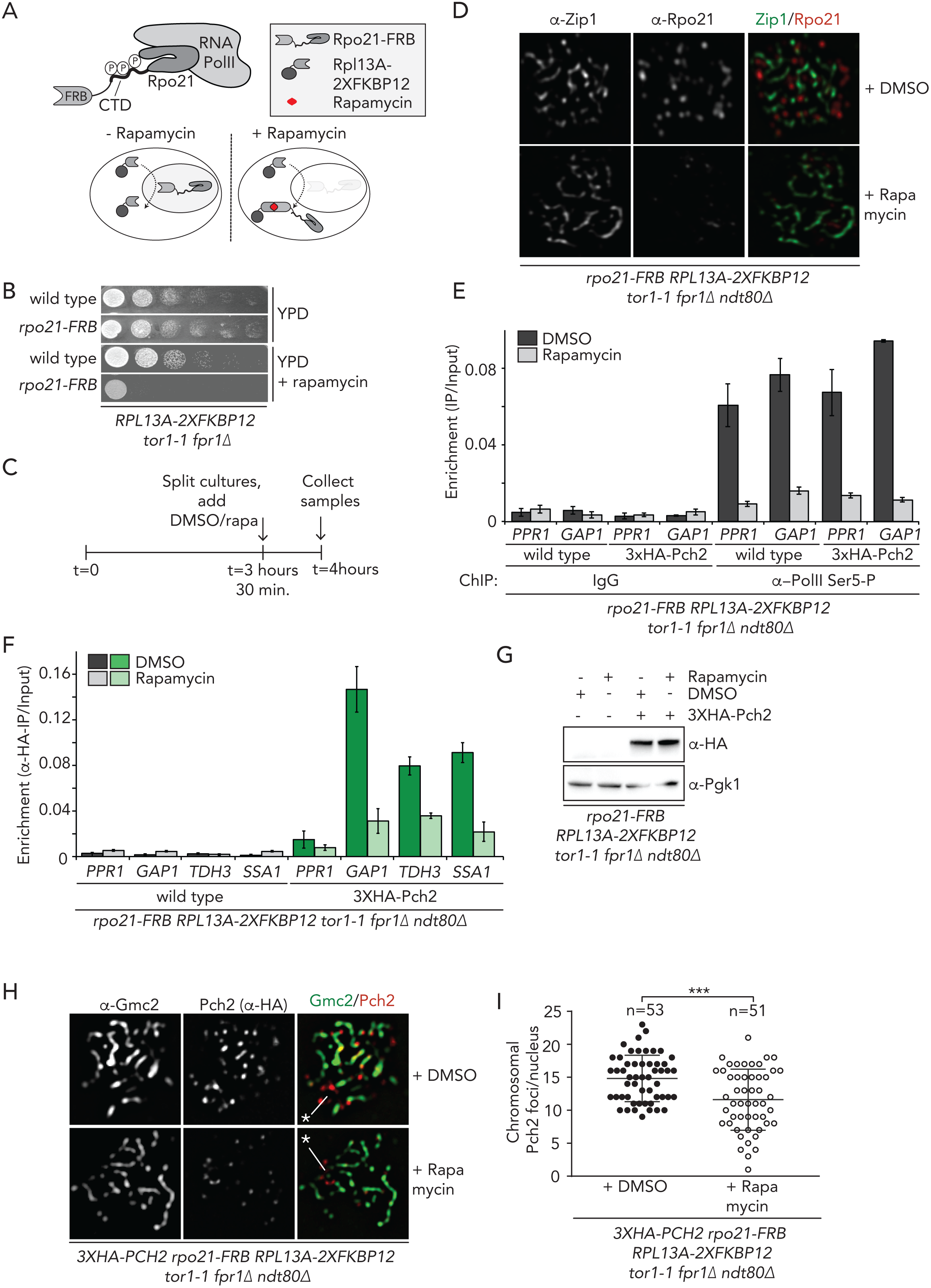
Transcription is required for recruitment of Pch2. A. Schematic of the anchor away system and Rpo21 within RNAPII. B. Dilution series of wild type and *rpo21-FRB* anchor away strains, grown on YPD or YPD + rapamycin solid medium. C. Schematic of treatment regimens used for anchor away experiments in D-H. D. Immunofluorescence of meiotic chromosome spreads in the *rpo21-FRB* anchor away treated with DMSO or rapamycin. E. ChIP-qPCR analysis of active transcription (α-phosphoserine 5 Rpo21 ChIP) in *rpo21-FRB* anchor away treated with DMSO or rapamycin at *PPR1* (yGV2390/yGV2391) and *GAP1* (yGV2597/yGV2598). Error bars represent standard error of at least three independent experiments performed in triplicate. F. ChIP-qPCR analysis of 3XHA-Pch2 in *rpo21-FRB* anchor away treated with DMSO or rapamycin at *PPR1* (yGV2390/yGV2391), *GAP1* (yGV2597/yGV2598), *HOP1*(yGV2607/yGV2608), *TDH3* (yGV2591/yGV2592), and *SSA1* (yGV2587/yGV2588). Error bars represent standard error of at least three independent experiments performed in triplicate. G. Expression analysis of 3XHA-Pch2 in *rpo21-FRB* anchor away treated with DMSO or rapamycin. H. Immunofluorescence of meiotic chromosome spreads in the 3XHA-Pch2 expressing *rpo21-FRB* anchor away cells, treated with DMSO or rapamycin. Chromosome synapsis was assessed by α-Gmc2 staining. I. Quantification of immunofluorescence as shown in H, treated with DMSO or rapamycin. *** indicates a significance of p≤ 0.001, Mann-Whitney U test.

**Figure 3.**
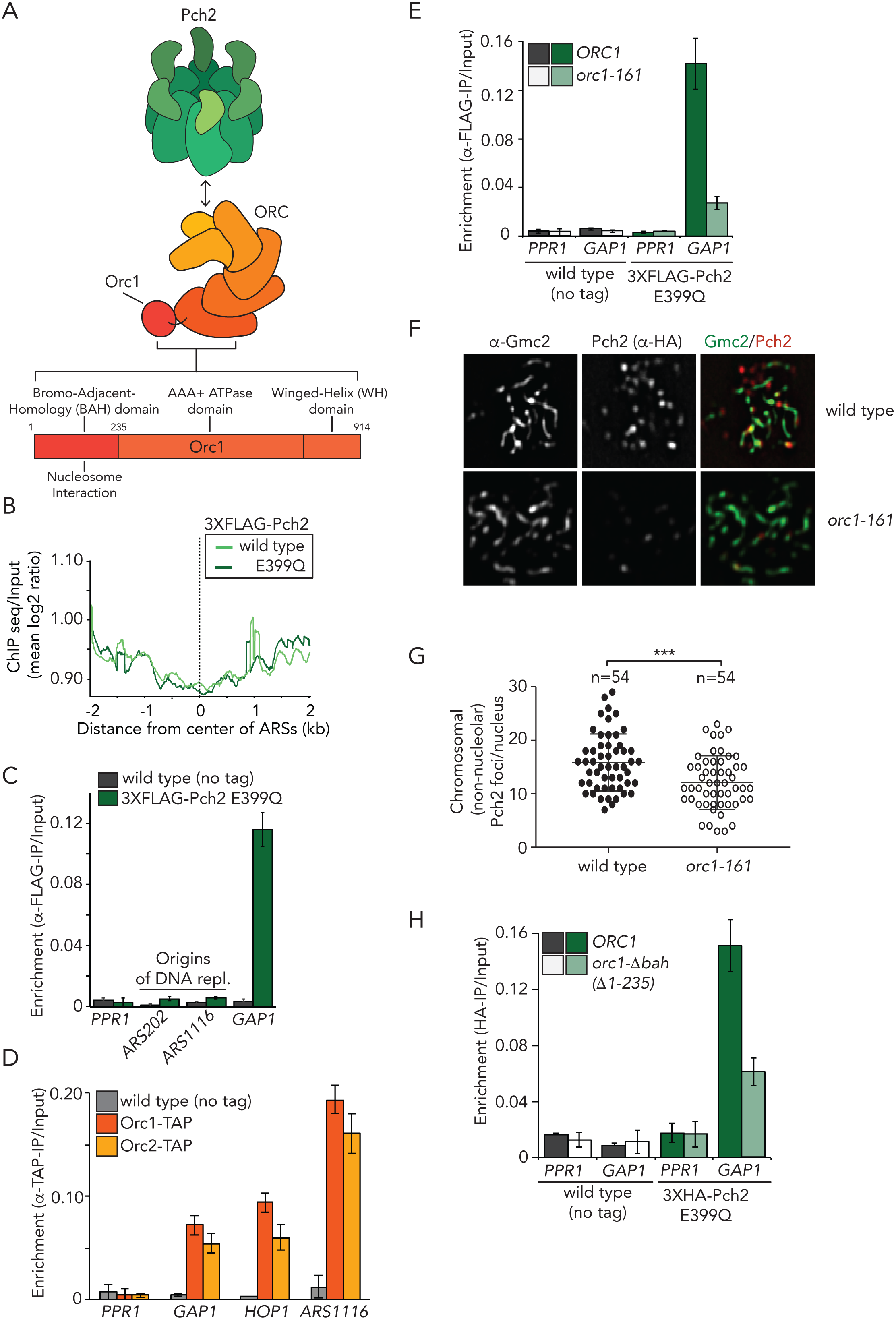
Interplay between Pch2, Orc1 and transcription. A. Schematic of Pch2 and ORC, including the domain organization of Orc1. B. Whole genome average plotting of 3XFLAG-Pch2 and 3XFLAG-Pch2-E399Q binding peaks (log2). Datasets were aligned relative to the center of ARSs. C. ChIP-qPCR analysis of 3XFLAG-Pch2-E399Q at *PPR1* (yGV2390/yGV2391), *GAP1* (yGV2597/yGV2598), *ARS202* (yGV2583/yGV2584) and *ARS1116* (yGV2577/yGV2578) during meiotic G2/prophase (4 hours). Error bars represent standard error of at least three independent experiments performed in triplicate. D. ChIP-qPCR analysis of Orc1-TAP and Orc2-TAP at *PPR1* (yGV2390/yGV2391), *GAP1* (yGV2597/yGV2598), *HOP1* (yGV2605/yGV2606), *ARS1116* (yGV2577/yGV2578) during meiotic G2/prophase (4 hours). Error bars represent standard error of at least three independent experiments performed in triplicate. E. ChIP-qPCR analysis of 3XFLAG-Pch2-E399Q in *ORC1* or *orc1-161* background at *PPR1* (yGV2390/yGV2391) and *GAP1* (yGV2597/yGV2598) during meiotic G2/prophase (4 hours). Experiments were performed at 23°C. Error bars represent standard error of at least three independent experiments performed in triplicate. F. Immunofluorescence of meiotic chromosome spreads in 3XHA-Pch2 expressing *ORC1* or *orc1-161* cells. Chromosome synapsis was assessed by α-Gmc2 staining. Experiments were performed at 23°C. G. Quantification of F. *** indicates a significance of p≤ 0.001, Mann-Whitney U test. H. ChIP-qPCR analysis of 3XHA-Pch2-E399Q in *ORC1* or *orc1Δbah* background at *PPR1* (yGV2390/yGV2391), *GAP1* (yGV2597/yGV2598) during meiotic G2/prophase (4 hours). Error bars represent standard error of at least three independent experiments performed in triplicate.

**Figure 4.**
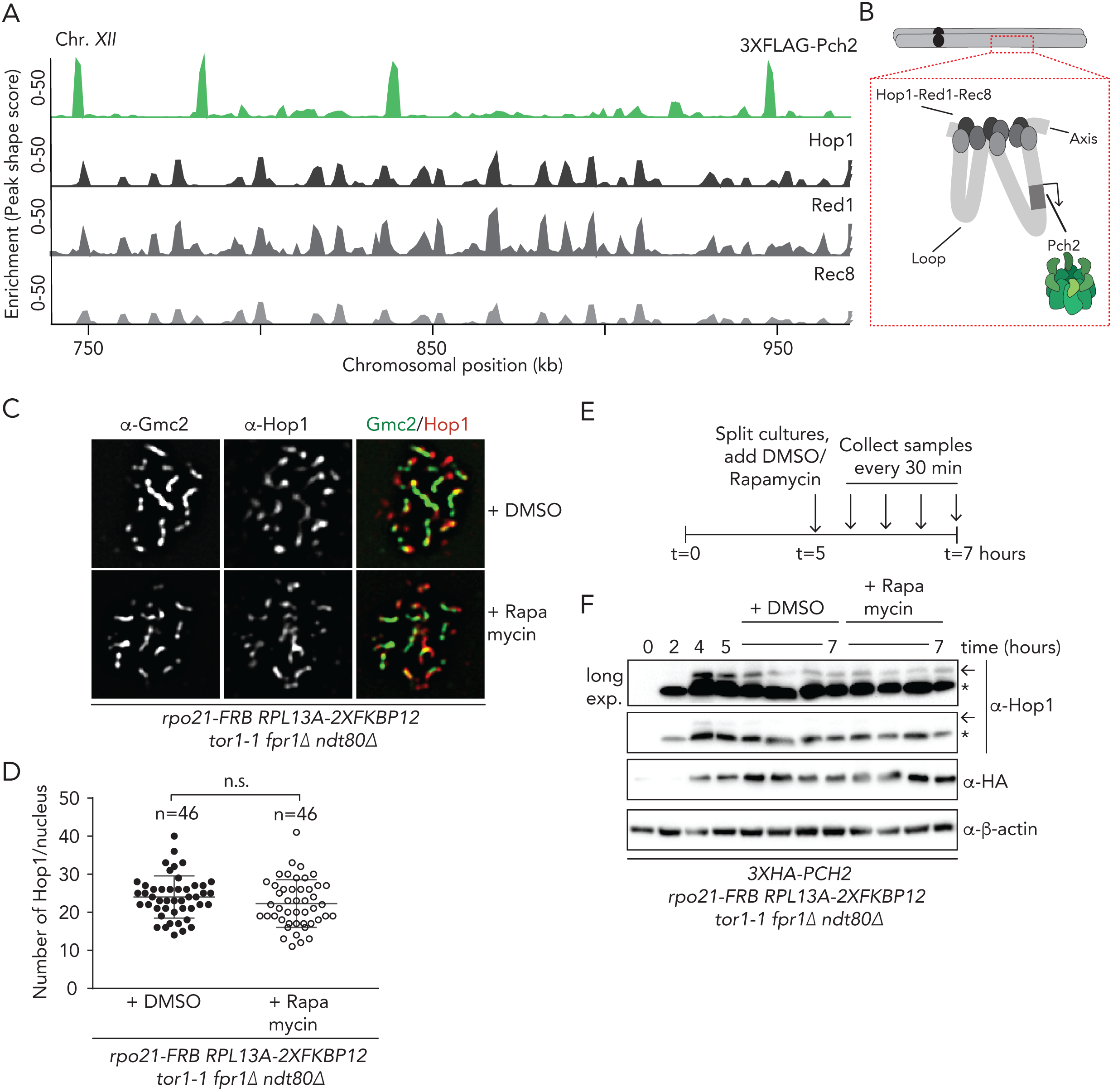
Functional analysis of the transcriptional-associated Pch2 chromosomal population. A. Representative image of ChIP-seq binding patterns for 3XFLAG-Pch2, Hop1, Red1 and Rec8. Data for Hop1, Red1 and Rec8 are from [8]. Shown is a region of Chromosome *XII* (chromosomal coordinates (kb) are indicated). B. Model depicting the proposed localization pattern of Pch2 on loops, within the meiotic chromosome loop-axis structure. C. Immunofluorescence of Hop1 on meiotic chromosome spreads in *rpo21-FRB* anchor away cells, treated with DMSO or rapamycin, using the regimen indicated in Figure 2C. Chromosome synapsis was assessed by α-Gmc2 staining. D. Quantification of F. n.s. (non-significant) indicates p>0.05, Mann-Whitney U test. E. Schematic of treatment regimens used for anchor away experiment, as shown in F. F. Western blot analysis of Hop1 and Pch2 (α-HA) in *rpo21-FRB* anchor away cells, treated with DMSO or rapamycin, as indicated in E. Upon DMSO or rapamycin treatment samples were taken every 30 minutes. Arrow indicates phosphorylated Hop1, * indicates non-phosphorylated Hop1.

**Figure 5.**
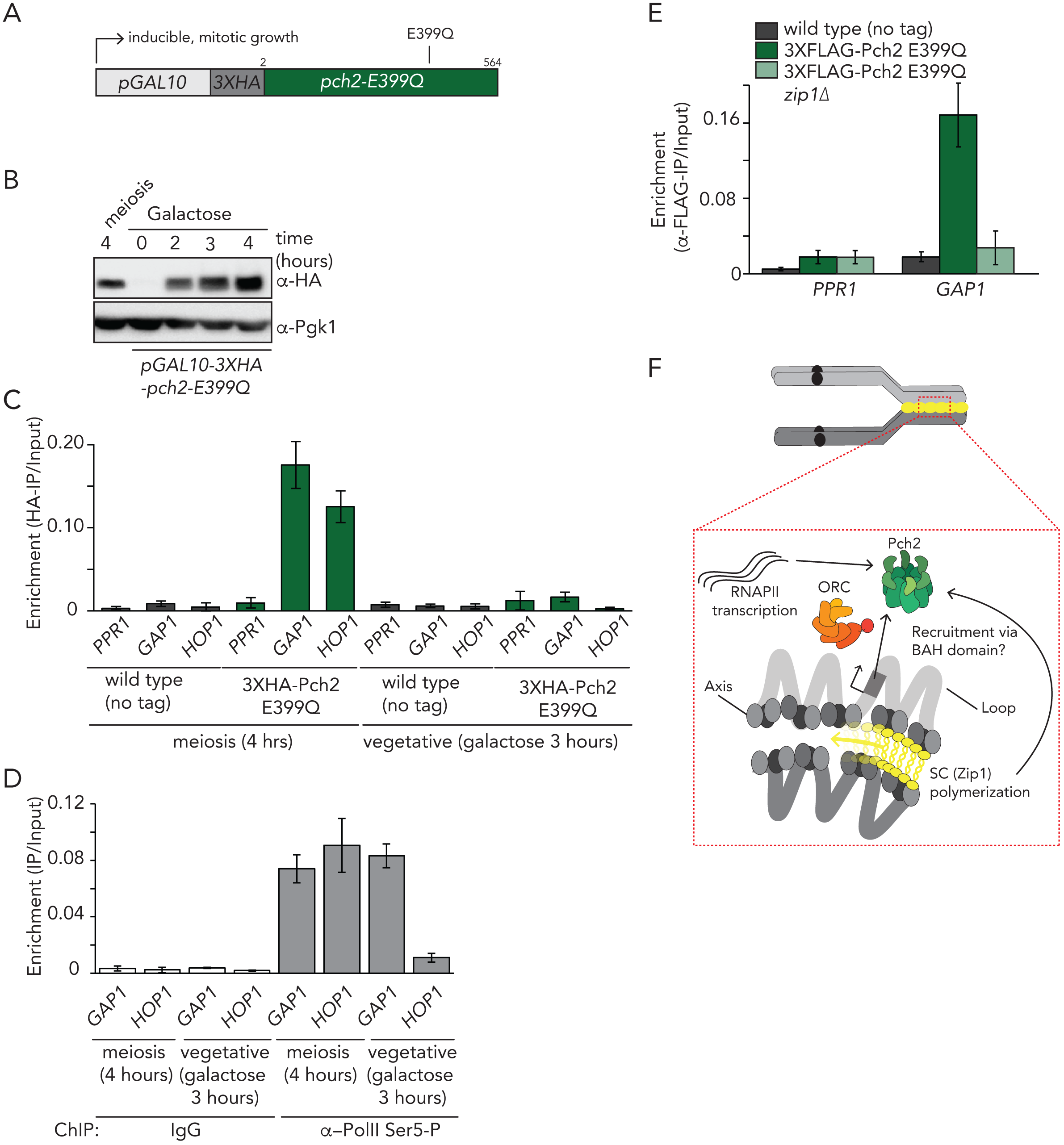
Requirements of Pch2 binding in mitosis and meiosis. A. Schematic of allele used for galactose-inducible mitotic expression of 3XHA-Pch2-E399Q. B. Western blot analysis of meiotic (*i.e.* endogenous) expression (4 hours) and ectopic expression of *pGAL10-3XHA-pch2-E399Q*. Hours of treatment with galactose are indicated. C. ChIP-qPCR analysis of 3XHA-Pch2-E399Q at *PPR1* (yGV2390/yGV2391), *GAP1* (yGV2597/yGV2598) and *HOP1* (yGV2605/yGV2606) during in meiosis and mitosis. Time (hours) is indicated. Error bars represents standard error of at least three independent experiments performed in triplicate. D. ChIP-qPCR analysis of active transcription (α-phosphoserine 5 Rpo21) *GAP1* (yGV2597/yGV2598), *HOP1* (yGV2605/yGV2606) in meiosis and mitosis. Hours are indicated. Error bars represent standard error of at least three independent experiments performed in triplicate. E. ChIP-qPCR analysis of 3XFLAG-Pch2-E399Q at *PPR1* (yGV2390/yGV2391) and *GAP1* (yGV2597/yGV2598) in wild type or *zip1Δ* cells during meiotic G2/prophase (4 hours). Error bars represent standard error of at least three independent experiments performed in triplicate. F. Model depicting the interplay between Pch2 binding, active transcription, Orc1 and chromosome organization.

### Nuclear depletion via the anchor-away method

Rpo21 was functionally depleted from the nucleus using the anchor away technique [32, 33]. Briefly, Rpo21 was tagged with FKBP12-rapamycin-binding (FRB), and this allele was introduced in strains harboring the anchor away background (*RPL13A-2xFKBP12 fpr1Δ tor1-1)* [32]. Nuclear depletion of Rpo21-FRB was achieved by addition of rapamycin at a concentration of 1 μM. Exact treatment regimens are indicated per experiment (see figure legends). For viability assays, serial dilutions of mitotically growing yeast cells were spotted on solid YPD containing 1 μM rapamycin for 2 days.

### Co-immunoprecipitation and western blots

100mL of SPO cultures (OD600 1.9), were harvested at 3000 rpm for 3 min at 4°C and washed once with ice-cold Tris-buffered saline (TBS) buffer (25 mM Tris–HCl, pH 7.4, 137 mM NaCl, 2.7 mM KCl). Cells were snap-frozen in liquid nitrogen and stored at −80°C until further use. Cells were resuspended in lysis buffer (50mM Tris-HCl pH 7.4, 150mM NaCl, 1% Triton X-100, and 1mM EDTA) containing protease inhibitors and broken with glass beads using bead beater (FastPrep-24, MP Biomedicals; 2 × 60 seconds at speed 6.0, incubated on ice in between for 5 min). Chromatin was sheared by sonication using a Bioruptor (Diagenode), 25 cycles of 30 seconds on/off, high power at 4°C. Lysates were clarified by centrifugation for 15 min at 16,000 x g at 4°C. Lysates were then immunoprecipitated with α-TAP antibodies using magnetic beads (Invitrogen), washed 4 times with buffer containing detergent and another time with the same buffer without detergent. Beads were eluted in 1X Loading buffer (50 mM Tris-HCl pH 6.8, 2% SDS, 10% glycerol, 1% β-mercaptoethanol, 12.5 mM EDTA, 0.02 % bromophenol blue) and the supernatant resolved by SDS-PAGE followed by Western blot. Protein extracts were prepared by using trichloroacetic acid (TCA) extraction protocol as previously described [28]. Samples were resolved by SDS-PAGE, transferred to nitrocellulose membranes and probed with the following primary antibodies diluted in 5% (w/v) nonfat-milk in TBS buffer + 0.1% Tween 20: α-Flag (Sigma-Aldrich, F3165), α-ORC2 (Abcam, 664906), α-HA (BioLegend, 901502), α-TAP (Thermo Scientific, CAB1001), α-Pgk1 (Invitrogen, 459250), α-Rpo21 (BioLegend, 664906), α-phosphoserine 5 Rpo21 (Thermo Scientific, MA518089), α-Histone-H3 (Abcam, AB1791), α-FRB (Enzo, ALX215-065-1), α-Zip1 (Santa Cruz Biotechnology, YC-19), α-Hop1 (kind gift of Nancy Hollingsworth, Stony Brook University, Stony Brook, USA), α-β-Actin (Abcam, AB170325), α-Histone H2A (Active Motif, AB2687477) or α-ORC (kind gift of Stephen Bell, MIT, Cambridge, USA). Membranes were incubated with horseradish peroxidase-conjugated goat anti-rabbit IgG, anti-mouse IgG and donkey anti-goat IgG (Santa Cruz Biotechnology). Proteins were detected with ECL (GE Healthcare) using a digital imaging system Image-Lab (Bio-Rad).

### Chromatin Immunoprecipitation (ChIP)

For ChIP experiments 100 mL SPO-cultures (OD600 1.9) were harvested 4 hours after entering meiosis, unless stated otherwise. Meiotic cultures or exponentially growing mitotic cultures were crosslinked with 1% methanol-free formaldehyde for 15 minutes at room temperature and the reaction was quenched with 125 mM Glycine. Cells were washed with ice-cold TBS, snap-frozen and stored at – 80°C. Cells were resuspended and broken with glass beads using a bead beater, as described above. Chromatin was sheared using either a Branson Sonifier 450 (microtip, power setting 2, 100% duty cycle, 3X for 15sec, 2 min on ice in between) or using a Diagenode Bioruptor UCD 200 (25 cycles of 30 seconds on/off, high power at 4°C). Cells were centrifuged at 13000 rpm for 10 min at 4°C. 10% of sample was removed for input. 550 µl of cell lysates were pre-incubated with the following antibodies for hours at 4°C prior to overnight incubation under rotation with magnetic Dynabeads-protein-G (Invitrogen): for ORC ChIP, 1 µl of α-ORC and 1 µl of isotype control antibody (α-rabbit IgG (Bethyl, P120-101). For TAP ChIP, 1 µl of α-TAP, and for HA ChIP, 1 µl of anti-HA. For RNA Pol-II ChIP, 1 µl of α-Rp021 or α-phosphoserine 5 Rpo21. Immunoprecipitates were incubated and washed as described above. For FLAG ChIP cells lysates were incubated with 30μl of 50% α-Flag-M2 affinity gel (Sigma-Aldrich, A2220) for 3 hours. Bound proteins were eluted using a 3XFLAG peptide (Sigma-Aldrich, F4799) as described in [34]. Subsequent steps (*i.e.* reversal of crosslinking, Proteinase-K and RNase-A treatments and final purifications and elutions) were performed as previously described in [35].

### ChIP-Seq library preparation

Preparation of paired-end sequencing libraries was performed using the Illumina TruSeq ChIP library preparation kit, according to the manufacturer’s guidelines. Ligation products were size-selected (250-300 bp) and purified from a 2% low-melting agarose gel using the MinElute Gel Extraction Kit (Qiagen). Ampure XP beads (Agilent) were used for cleanup steps and size selection. The final purified product was quantitated using Picogreen in a QuantiFluor dsDNA System (Promega). Sequencing was performed on the Illumina NextSeq 500 platform at the Max Planck Genome Centre Cologne, Germany.

### Processing of ChIP seq data

Preliminary quality control of raw reads was performed with FastQC. Illumina raw sequences were then filtered for removal of low quality and duplicated reads, adapters and low-quality bases using the SAM tools. Paired-sequencing reads from three biological independent replicas (ChIPed DNA and their respective inputs from 3XFLAG-Pch2 and 3XFLAG-tagged-Pch2-*E399Q* expressing cells) were aligned and mapped to the S288C (SacCer3) genome assembly with CLC Genomics Workbench 10, allowing for maximal 2 mismatches in 100bp. Duplicated reads were removed. The resulted mappings were then used as input for peak calling. The peaks shown were obtained through normalization with inputs. In order to obtain high-quality peaks, a p-value of 1e-15 was used for calling the peaks. Peak calling was performed with CLC-genomics. The CLC peak caller is based on a Gaussian filter that derives the mean variance from cross-correlations obtained from the ChIP-seq data and its respective input control, thus avoiding detection of peaks from regions with strong sequencing biases or potential PCR artefacts originated during library preparation [36]. Bowtie reading maps was also used as input for peak calling and resulted in similar peak profiles. All peaks were manually verified prior to analysis aiming to exclude those which were not present in at least two of the three technical replicates. For RNA-seq data analyses shown in Supplementary Figure 1C and D, reads obtained from (GSE131994, 3 hours in meiosis) [23], were mapped to the SK1 genome background (SGRP) and analyzed using the RNA-seq platform on CLC-genomic workbench. For analysis shown in figure Supplementary Figure 2A, peak calling was performed by subtracting NLS-GFP ChIP-seq data (SRR2029413) [37] from the Pch2 ChIP-seq dataset. Hop1 (GSM1695721 and GSM165720), Red1 (GSM1695718 and GSM1695716) and Rec8 (GSM1695724 and GSM1695722) ChIP-seq datasets were from [8]. For Supplementary Figure 6F, Hop1 ChIP-seq datasets (wild type; GSM2818425 and GSM2818423, *pch2Δ*; GSM2818432 and GSM2818436) were used from [38].

### Computational analyses

Log2 ratios (IP/Input) of the genome-wide enrichment of ChIP-seq (3XFLAG-Pch2 and 3XFLAG-tagged-Pch2-*E399Q*) was generated by MACs peak calling using the Read Count Quantitation algorithm of Seqmonk (version 0.34.1). To generate distance maps, <u>O</u>pen <u>R</u>eading <u>F</u>rames (ORFs) and tRNAs, (gene annotations and chromosomal coordinates were obtained from SGD (http://www.yeastgenome.org)) and <u>A</u>utonomously <u>R</u>eplicating <u>S</u>equences (ARSs) (coordinates and sequences for the 798 (confirmed, likely, and dubious) origins were downloaded from the OriDB (version 2012 (http://www.oridb.org)) were transformed into BED files using the table browser tool (USCS) [39]. Intensity was parsed into 100 base pair bins and Pch2 binding sites were aligned at the center of ORFs, tRNAs and ARSs. Venn diagrams were generated using the web-based Venn diagram generator from http://jura.wi.mit.edu/bioc/tools/venn.php. Pearson correlation was performed with datasets from the CLC-genomics peak shape score for each Pch2 binding gene and their specific expression values obtained from analysis of RNA-seq data [23] (Supplementary Figure 1D). To determine whether the Pch2 binding genes were enriched in certain functional categories, gene ontology analysis was conducted using the SGD GO term finder (molecular function) (https://www.yeastgenome.org/goTermFinder) at a p-value cutoff of 0,01.

### ChIP-qPCR

ChIP and Input samples were quantified by qPCR on a 7500 FAST Real Time PCR machine (Applied Biosystems). The percentage of ChIP relative to input was calculated for the target loci as well as for the negative controls. Enrichment [relative to time untagged control or IgG (control)] was calculated using the ΔCt method as follows: 1/(2^[Ct−Ctcontrol]). Primers sequences (including primer efficiency) covering the various loci are listed in the Supplementary Data.

### RNA extraction and RT-qPCR

For RNA extraction, 15 mL meiotic cultures were harvested and total RNA was extracted using the hot-acidic phenol method [40], with some minor modifications. Cells were resuspended in 600 µl of freshly prepared TES buffer (10 mM Tris-Cl, pH 7.5 10 mM EDTA 0.5% SDS). 600 µl of acidic-phenol (Ambion) was added and the solution was immediately vortexed vigorously for 30 seconds. Samples were incubated at 65°C for 90 min under rotation at 300 rpm. The solution was kept on ice for 10 minutes and spinned down at 14000 rpm for 10 minutes at 4°C. The aqueous top layer was transferred to a new tube and 600 µl of clorophorm was added and immediately vortexed. Cells were centrifuged as above after which the aqueous layer was transferred to a new pre-chilled eppendorf tube. RNA was precipitated overnight at −20°C with 2.5 volumes of 100% ethanol and 10% (v/v) sodium acetate, pH 5.4 and washed with 75% ethanol. After drying on ice, RNA was eluted with RNase free water and stored at −80°C. cDNA was generated using the superscript-III reverse transcriptase (Invitrogen) according to the manufacturer’s protocol. Briefly, 1-2 µg of total RNA was used in a 20 µl reaction mixture using random primer mix or oligodT-20 (Invitrogen). Relative amounts of cDNAs of various genes were measured by real time quantitative PCR (RT-qPCR). Expression of every gene was normalized to expression of *18S* (RNA Pol I transcript) or β-Actin (*ACT1*) from the same RNA preparation. Oligo sequences are available in Supplementary Data.

### Chromosome spreads and immunofluorescence

Chromosome spreading was performed as described in [14]. For immune staining, the following antibodies were used: α-HA (Roche, 11867423001), α-Zip1 (Santa Cruz Biotechnology, YN-16), α-Gmc2 (a kind gift of Amy MacQueen, Wesleyan University, Middletown, CT, USA), α-Rp021 and α-Hop1 (home made). α-Hop1 were raised against full length 6-His-tagged Hop1 expressed in bacteria. Hop1 was purified via affinity purification followed by ion-exchange chromatography, and used for immunization. Antibody production was performed at the antibody facility of the Max Planck Institute of Molecular Cell Biology and Genetics (Dresden, Germany). DNA was stained with 4’,6-Diamidine-2’-phenylindole dihydrochloride (DAPI). Images were obtained using a DeltaVision imaging system (GE Healthcare) using a sCMOS camera (PCO Edge 5.5) and 100x 1.42NA Plan Apo N UIS2 objective (Olympus). Deconvolved images (SoftWoRx software 6.1.l and/or z-projected) using the SoftWoRx software 6.1.l). were quantitated using Imaris Software (Oxford Instruments).

## Results

We aimed to generate a detailed genome-wide mapping of the chromosomal localization pattern of Pch2, using chromatin immunoprecipitation followed by deep sequencing (ChIP-seq). For this, we employed a NH2-terminal 3xFLAG-tagged wild type version of Pch2, and a mutant allele harboring an E>Q substitution at position 399 within the AAA+ Walker-B motif (*pch2-E399Q*) (Figure 1A). This mutant is expected to cause impaired ATP hydrolysis, and equivalent mutations have been used to stabilize interactions between AAA+ proteins and clients and/or adaptors [41]. We anticipated that Pch2-E399Q would exhibit increased association to chromosomal regions as compared to its wild type counterpart, which could aid in revealing details regarding Pch2 recruitment and/or function. Strains expressing wild type Pch2 and Pch2-E399Q (Figure 1B) were used to generate ChIP-seq datasets during meiotic G2/prophase. We compared independent ChIP-seq datasets for wild-type and *E399Q* Pch2 (performed in triplicates in both cases) and found that these datasets exhibited highly correlated distributions, both at a genome-wide level and at individual loci. Distribution patterns were highly reproducible among replicas, and we averaged maps from three independent experiments for Pch2 wild-type and E399Q for further analyses. In addition, for several follow-up experiments (see below), we used both alleles (*i.e.* wild type and E399Q) interchangeably. Our genome-wide ChIP analyses revealed that Pch2 is recruited to distinct regions on all 16 budding yeast chromosomes (Supplementary Table 1). The use of a peak-calling algorithm with a stringent threshold [36] revealed that the vast majority (*i.e.* ∼97 %; for wild type Pch2 434 out of 447 peaks, for Pch2-E399Q 525 out of 540 total peaks) of the identified peaks (p-value: 1e-15, see material and methods) localized within the coding sequences (CDS) of a subset of RNAPII-transcribed genes (see Figure 1C and 1D for typical examples of binding patterns, Supplementary Figure 1A). ∼3% of peaks represent binding sites with relatively low peak shape scores at centromeres and some telomeric regions. We did not observe Pch2-association within promoters (*i.e.* directly upstream of the transcriptional start sites (TSSs)) of these Pch2-bound genes. Pch2 peaks were evenly distributed throughout CDSs and located downstream of TSSs, showing symmetrical ChIP patterns that are reminiscent of ChIP signals for transcriptionally-engaged RNAPII (Figure 1E). We plotted the average global difference between Pch2 wild-type and Pch2 *E399Q* for the 434 common peaks. Association of Pch2 E399Q to the common binding sites (434 peaks) is stronger relative to wild-type Pch2, as judged by the differences in peak shape score (Supplementary Figure 1B). In general, individual Pch2 E399Q peaks had either similar or higher peak shape scores in comparison to wild-type Pch2. Based on the biochemical characteristics of AAA+ enzymes, the Pch2-E399Q is expected to exhibit stronger binding to clients and/or adaptors [41], and these increased binding patterns thus suggest that the observed binding sites represent biochemically meaningful interactions. In addition to the observed association of Pch2 with RNAPII-transcribed genes, we also found evidence for specific Pch2-binding patterns within the rDNA array (RCS and GV, unpublished observations). These binding patterns might relate to the observed enrichment of Pch2 within the nucleolus [17, 28]. Here, we focus on the Pch2 binding patterns across the non-rDNA, euchromatic part of the genome. We queried sequences of the 434 genes for motifs using MEME motif-finding software (Bayley et al, 2009), but this did not identify motifs with any degree of significance (data not shown), suggesting that recruitment of Pch2 is not directly connected with any obvious DNA sequence-directed factors. We next performed a search for enriched Gene Ontology (GO) terms showing a representation for genes involved in various metabolic processes (Supplementary Table S2). In addition to these GO terms, Pch2-association was also enriched within sporulation-induced (*i.e.* meiosis-specific) genes. Analysis of genome-wide transcriptome (*i.e.* total RNA-seq) datasets from cells that synchronously progress through meiosis [23] showed that all of the genes occupied by Pch2 are transcribed during meiosis, suggesting that transcription is involved in the recruitment of Pch2 to these CDSs. To test if transcriptional “strength” of defined genes was predictive of Pch2 binding, we stratified the transcribed genes from the RNA-seq dataset into high, medium and low expression strength (following previously established procedures [42]), and compared expression strength of Pch2-associated genes with these bins (Supplementary Figure 1C). This analysis showed that Pch2-associated genes produce average mRNA levels, with a wide distribution. We detected a weak correlation between the peak shape score of individual Pch2-binding sites and the expression level of the corresponding CDS (Pearson’s test, R^2^=0.0026, Supplementary Figure 1D). This indicates that, although Pch2 associates with actively transcribed genes, expression strength is not a determining factor for Pch2 binding. Underscoring this interpretation is the fact that many highly expressed genes do not show significant Pch2 enrichment peaks. ChIP analysis can be plagued by artefactual ChIP-enrichments, some of which have been observed within highly-expressed, RNAPII-transcribed genes and within non-coding, but equally highly-expressed RNAPIII-transcribed tRNAs (these regions are often referred to as hyperChIPable regions [37]). We performed several analysis and experiments to exclude artefactual binding effects in our ChIP datasets, and we discuss these in the Supplementary Results (see also Supplementary Figure 2). Based on these experiments and on additional experiments that are described below, we are confident that our Pch2 datasets inform on physiologically relevant biological behavior.

To further explore the information gathered by our ChIP-seq results, we employed ChIP followed by real-time quantitative PCR (ChIP-qPCR). We designed oligo’s flanking regions spanning three different portions of a selected Pch2 binding gene, *GAP1. PPR1*, an RNAPII-transcribed gene to which Pch2 showed no association by ChIP-seq, was used as a negative control. 3xFLAG-Pch2-*E399Q* associated to the three regions spanning the *GAP1* gene body to similar extents (Figure 1F), whereas no significant association was observed at the *PPR1* locus (Supplementary Figure 1E). We confirmed transcriptional activity at *GAP1* and *PPR1* by ChIP analysis of active RNAPII (as judged by ChIP-qPCR analysis of phosphorylated C-terminus domain of Rpo21 (α−PolII-Ser5-P ChIP) (Supplementary Figure 1F). The C-terminal domain (CTD) of Rpo21 harbors a series of YSPTSPS heptad repeats that are hyperphosphorylated during transcription (reviewed in [43]). Phosphorylation of Rpo21-Serine 5 can be used as a read-out to assess active engagement of RNAPII during transcription elongation. We note that *GAP1*, a gene highly expressed both in exponentially cycling cells and during meiosis was also identified as a site of hyperChIPpability (*i.e.* it is one of the 19 genes that shows overlap between our Pch2 datasets and the NLS-GFP dataset). Importantly, in agreement with the results obtained by ChIP-seq, we confirmed the association of wild-type Pch2 and Pch2-E399Q to three additional binding genes tested (*HOP1, TDH3* and *SSA1*), none of which were present in the hyperChIPable dataset (Figure 1G and H). Based on this (and on several additional lines of evidence; see Supplementary results and discussion), we are confident that Pch2 enrichment at *GAP1* represents a true enrichment (and that *GAP1* can be used to further investigate the biology behind Pch2 recruitment). Using ChIP-qPCR we confirmed increased binding of Pch2-E399Q as compared to wild-type Pch2 (Figure 1G and H). In addition to its catalytic AAA+ domain, Pch2 also possess a non-catalytic NH2-terminal domain (NTD) (Figure 1A) [19]. The NTDs of AAA+ ATPases are required to allow AAA+ proteins to interact with clients and adaptors [19, 41]. Removal of the NTD of Pch2 abrogated the association of Pch2 to individual selected genes, indicating that the NTD is required for recruitment of Pch2 to gene bodies (Figure 1G and H). In total, we conclude that during meiotic G2/prophase, Pch2 associates within the body of a selected group of RNAPII-associated genes, and that recruitment depends on specific characteristics of AAA+ proteins.

We next wanted to establish whether global inhibition of RNAPII transcription affected Pch2 occupancy. To inhibit RNAPII-dependent transcription, we initially used 1,10-Phenanthroline − a small molecule that most likely acts as a magnesium chelator −, which has previously been described to inhibit RNAPII-dependent transcription [30]. Meiotic yeast cultures expressing 3XFLAG-Pch2 E399Q were treated with 1,10-Phenantroline for 1 hour. Under these conditions, we observed a substantial effect on *GAP1* mRNA levels, in cells treated with 1,10-Phenantroline, whereas Pch2 protein levels were not affected (Supplementary Figure 3A-C). Inhibition of global transcription reduced Pch2 association to *GAP1* gene by ∼50% compared to mock control (Supplementary Figure 3D), consistent with a role for transcription in promoting the recruitment/association of Pch2 to regions of active transcription. To achieve a more complete and specific inhibition of RNAPII, we employed the anchor-away technique [32], which has been used to successfully deplete chromosomal proteins during meiosis [14, 23, 38, 44]. This technique is based on an inducible dimerization system that rapidly depletes nuclear proteins based on ribosomal flux, with the aid of a tagged anchor-protein, Rpl13A (Rpl13a-2XFKBP12). Rapamycin induces the formation of a ternary complex with a protein of interest that is tagged with FRB (<u>F</u>KBP12-<u>R</u>apamycin <u>B</u>inding-FRB domain of human mTOR) (Figure 2A). Successful inhibition of RNAPII using the same genetic approach has been described in vegetative cells [33], and we tagged the largest subunit of RNAPII (Rpo21) with the FRB tag (Supplementary Figure 3E). As expected, *rpo21-FRB* cells exhibited severe growth defects in the presence of rapamycin (Figure 2B). Immunofluorescence of Rpo21 on meiotic chromosome spreads after exposure with Rapamycin for 30 minutes demonstrated efficient nuclear depletion of Rpo21-FRB during meiosis (Figure 2C and D). We additionally performed ChIP-qPCR analysis, using antibodies against phosphorylated-Serine 5 of Rpo21 in meiotic cells following addition of rapamycin or DMSO. Our ChIP shows that the relative occupancy of RNAPII within *PPR1* and *GAP1* coding regions was substantially reduced after 30 minutes rapamycin treatment in cells expressing Rpo21-FRB (Figure 2E), further demonstrating that RNAPII is depleted from the nucleus under this treatment regimen. Importantly, ChIP analysis revealed that Pch2 binding was significantly reduced in Rpo21-FRB-tagged strains treated with rapamycin for 30 minutes (Figure 2F), whereas the protein levels of Pch2 were unaffected (Figure 2G). We conclude that active transcription is required to promote the recruitment/association of Pch2 to a defined subset of transcriptionally active genes. We next addressed whether a reduction of Pch2 binding to these regions upon transcriptional inhibition could be corroborated through independent, cytological methods. For this, we performed immunofluorescence on spread chromosomes to quantify the chromosomal association of Pch2 during meiotic G2/prophase, and found that a brief inhibition (*i.e.* 30 minutes) of active transcription (via Rpo21-FRB nuclear depletion) triggered a significant reduction of Pch2 chromosome-associated foci within synapsed chromosome regions (as identified by staining with the SC component Gcm2 [45]) (Figure 2H and I). We note that the loss of Pch2 from synapsed chromosomes under these conditions is partial, which is in contrast to the complete loss of Pch2 from chromosomes that is seen in *zip1Δ* cells [17]. This difference might indicate an incomplete inhibition of transcription-dependent recruitment under the used conditions. Alternatively, it could suggest the presence of additional chromosome-associated pools of Pch2, that are recruited independently of transcription. Importantly, the nucleolar pool of Pch2 (identified by typical structure and lack of association with SC structures) appeared unaffected by RNAPII inhibition, suggesting that nucleolar recruitment does not depend on RNAPII-dependent transcription (Figure 2H). Together, these data identify active RNAPII-dependent transcription as a factor that contributes to the recruitment of Pch2 to euchromatic chromosome regions during meiotic G2/prophase.

We aimed to better understand the recruitment of Pch2 to chromosomes and its connection with transcription. For this, we focused on Orc1, a factor involved in Pch2 recruitment to the nucleolus [28, 46]. Orc1 is a component of the Origin Recognition Complex (ORC), a six subunit (Orc1-6) hexameric AAA+ ATPase (Figure 3A), and we recently showed that Pch2 directly interacts with ORC [47]. The first step in DNA replication occurs when the ORC recognizes and directly binds hundreds of origins of replication (also knowns as autonomously replicating sequences (ARSs)) across the genome (reviewed in [29]). Given the direct interaction between ORC and Pch2 and the well-established association of ORC with origins, we first queried whether Pch2 was associated with origins. We plotted Pch2 occupancy (log2 genome-wide enrichment of Pch2 wild-type and E399Q) around the center of 798 predicted ARSs (from OriDB (http://cerevisiae.oridb.org); this list includes dubious, likely and confirmed ARSs). This analysis did not reveal enrichment for Pch2 at or around ARSs (Figure 3B). To confirm this observation, we recalled Pch2 peaks using a substantially reduced significance threshold (*i.e.* a p-value of 0.5 instead of the previously mentioned 1e-15) [36], but this equally did not show any enrichment (data not shown). Further, using algorithms previously employed to detect association of ORC with ARSs (*i.e.* MACs peak calling [48]) similarly failed to reveal enrichment of Pch2 to ARSs (data not shown). The observation that Pch2 does not significantly associate with origins of replication was further strengthened by ChIP-qPCR investigating individual origins (Figure 3C). We conclude that Pch2 is not detectably associated with euchromatic origins of replication during meiotic G2/prophase, hinting that the interaction between Pch2 and Orc1 occurs away from origins of replication [47]. Previous studies have described association of components of the replication machinery (including ORC) to actively transcribed, protein coding genes [49-51], and we thus considered the possibility that ORC exhibited a binding pattern similar to that of Pch2 during meiosis. We analyzed Orc1-TAP binding by ChIP-qPCR: as a positive control, we measured its association to *ARS1116*, and as expected found Orc1 to be highly enriched at this site (Figure 3D). Strikingly, we also detected substantial binding of Orc1-TAP with *GAP1* and *HOP1*, but not *PPR1 (*Figure 3D and Supplementary Figure 4A). Similar results were seen for Orc2-TAP (Figure 3D), suggesting that the ORC complex may be associated to (non-origin) genomic regions that are also occupied by Pch2. To further explore the possibility that Orc1 functions upstream of Pch2 with respect to its localization − as has been suggested within the nucleolus/rDNA [28, 46] − we made use of a temperature sensitive allele of *ORC1, orc1-161* (Supplementary figure 4B). Pre-meiotic DNA replication is delayed for ∼1 hour in *orc1-161* cells, but meiotic progression is otherwise normal (Supplementary figure 4C) [28]. This mutation severely diminished ORC association with origins of replication (as measured by ORC2-TAP ChIP; Supplementary figure 4D), even at permissive temperatures (23°C), likely due to a reduction in Orc1 protein levels (Supplementary figure 4B) [52]. The levels of Orc2-TAP at *GAP1* were reduced under these conditions (Supplementary figure 4D). We next performed Pch2 ChIP-qPCR in yeast strains harboring temperature-sensitive alleles of either *ORC1 (orc1-161)* or *ORC2 (orc2-1)*. Strikingly, as shown in Figure 3E, in *orc1-161* cells, Pch2 levels were strongly depleted at *GAP1*, indicating that Orc1 contributes to the chromosomal association of Pch2, also outside the nucleolus. Pch2 protein levels were unchanged under these conditions (Supplementary Figure 4E). Orc2 was readily detected in *ORC2* cells but barely detected in *orc2-1* cells (Supplementary figure 4F). In contrast to the effects seen in *orc1-161*, ChIP-qPCR analysis of 3xHA-Pch2-E399Q revealed no effect of *orc2-1* on recruitment to GAP1 (for these experiments, the cultures were shifted to 30°C once the meiotic program was induced) (Supplementary Figure 4G and H). Thus, although we cannot rule out that incomplete inactivation of Orc2 in *orc2-1* obscures effects on Pch2 recruitment, these data collectively strongly suggest that Orc1 is involved in promoting the chromosomal association of Pch2. Cytological analysis showed that Pch2 localization to the nucleolus/rDNA is severely impaired in *orc1-161* cells [28]. Our ChIP analysis indicated that Orc1 also contributes to recruitment at non-rDNA loci, and accordingly, using immunofluorescence on spread chromosomes we found that in *orc1-161* cells Pch2 localization was diminished within non-rDNA regions (Figure 3F and G). Together, these analyses indicate that ORC, and particularly Orc1, are involved in and affect the localization of Pch2 at euchromatic chromosomal regions. Orc1 contains a <u>B</u>romo <u>A</u>djacent <u>H</u>omology (BAH) domain, a nucleosome-binding domain [53] that contributes to ORC’s ability to bind origins of replication [54] (Figure 3A). We and others have revealed a role for the BAH of Orc1 in controlling rDNA-associated functions [28, 55], and we interrogated whether occupancy of Pch2 to regions of transcriptional activity was affected in *orc1Δbah* cells. Indeed, deletion of the BAH domain of Orc1 significantly reduced the association of Pch2 to sites of active transcription (Figure 3H and Supplementary Figure 4I and J). Collectively, these results suggest that ORC/Orc1 may use its nucleosome-binding capacity (endowed by the BAH domain of Orc1) to bind to and recruit Pch2 to non-canonical (*i.e.* non-origin) genomic loci that are defined by transcriptional activity. In light of this, it is interesting to note that Orc1-BAH binding to nucleosomes − in contrast to a related BAH domain in Sir3 [56] − is insensitive to the acetylation state of Histone H4, which could allow effective engagement with nucleosomes within euchromatin [55].

Hop1, a HORMA-domain containing client of Pch2 is a central component of the meiotic axis structure [7, 14, 17, 24-27, 57]. Zip1-dependent SC assembly (which drives Pch2 recruitment [17]), is established on the axial element of the meiotic chromosome structure, and Hop1 and Zip1 are therefore expected to reside in molecular proximity of each other (at and near chromosome axis sites, respectively). As such, one hypothesis is that Pch2 is also enriched at meiotic axis-proximal sites. We compared the binding patterns of Pch2 to those of axial components (*i.e.* Red1, Hop1 and Rec8) by plotting available ChIP-seq datasets for Hop1, Red1 and Rec8 [8] and overlaying them with our Pch2 ChIP-seq datasets (Figure 4A). Hop1, Red1 and Rec8 showed highly similar binding patterns [7] [8], but strikingly, the binding patterns of Pch2 qualitatively diverged from the patterns of these axial elements (Figure 4A, and Supplementary Figure 5A and B). Specifically, Pch2 patterns did not show the similar frequency along chromosomal regions, and showed little overlap with the binding patterns of meiotic axis-factors. We conclude that the pool of Pch2 that we identified does not co-localize with the meiotic axis factors Hop1, Red1 and Rec8, and propose that, within the loop-axis organization of meiotic chromosomes, Pch2 associates with (a selected group of) genes that are located within loops that emanate away from the Hop1-Red1-Rec8-defined axis (Figure 4B). Because of this apparent spatial separation between Pch2 and its axial substrate Hop1, we asked whether impairing the recruitment of Pch2 to regions of active transcription affected the chromosomal abundance of Hop1 in meiotic G2/prophase. In *pch2Δ* cells, abundance of Hop1 on synapsed chromosomes is increased [14, 17, 58]. In addition, phosphorylation of Hop1 [59] increases in *pch2Δ* cells [14]. We used cytological and ChIP-based approaches to test effects of acute removal of Pch2 from regions of transcriptional activity (via our Rpo21-FRB-based system) on Hop1 chromosomal abundance. First, we quantified Hop1 chromosomal association with synapsed chromosomes under conditions where RNAPII-inhibition caused diminished localization of Pch2 (*i.e.* 30 minutes long exposures to rapamycin) (Figure 4C and D). Under these conditions, we did not observe significant changes in Hop1. We exposed cells to rapamycin for longer periods (*i.e.* 90 instead of 30 minutes), and under these conditions, Pch2 loss from chromosomes was more pronounced as compared to the loss observed upon short treatments (Supplementary Figure 6A-C). However, this was not accompanied by a detectable increase in the abundance of Hop1 on synapsed chromosomes (Supplementary Figure 6D-E). These experiments were all performed in early meiotic G2/prophase (*i.e.* after 4 hours into the meiotic program), and we also investigated effects later in meiotic G2/prophase (*i.e.* after 8 hours (in *ndt80Δ* cells, to prevent exit from G2/prophase). Also under these conditions, brief inhibition of RNAPII did not lead to significant effects on Hop1 chromosomal recruitment (data not shown). We next investigated Hop1 chromosomal levels by ChIP-qPCR, based on published Hop1 ChIP-seq datasets in wild type and *pch2Δ* cells [38], but equally did not find effects of acute inhibition of RNAPII on Hop1 chromosomal abundance at a selected locus (Supplementary Figure 6F and G). Finally, we probed the accumulation of the phosphorylated version of Hop1 (identified by a slower migrating band on SDS-PAGE gels). In accordance with earlier observations, we did not observe differences in the amount of phosphorylated Hop1 under conditions of RNAPII depletion (Figure 4E and F). Based on all cumulative experiments, we suggest that the pool of Pch2 that is associated with sites of active RNAPII-dependent transcription is not involved in regulating the chromosomal association of Hop1. This conclusion is in agreement with the observation that the binding sites for Pch2 that we identify here are not overlapping with chromosome axis sites, as defined by Hop1 (and Red1/Rec8).

Finally, we aimed to address further requirements for Pch2 association with selected regions of active RNAPII-dependent transcription. Most of the identified binding regions fall within genes that are also active in vegetatively growing cells (with the exception of a subset of meiosis-specific genes). In addition, Orc1/ORC is equally present in vegetatively growing cells. Therefore, we investigated whether ectopic expression of Pch2 − normally only expressed in meiosis − was sufficient to promote association with selected regions of binding, as identified in our ChIP analysis. We generated a galactose-inducible allele of Pch2 (*pGAL10-3HA-pch2-E399Q*) to induce Pch2-E399Q to protein levels comparable to those observed in meiotic G2/prophase (Figure 5A and B). ChIP-qPCR analysis revealed that, in contrast to the situation in meiotic cells, Pch2-E399Q was unable to associate with *GAP1* in mitosis, despite the fact that this gene was actively transcribed during vegetative growth (Figure 5C and D). These results show that active transcription and presence of Orc1/ORC are not sufficient for the association of Pch2 with selected regions of active transcription, and instead suggest the presence of meiosis-specific factors that ‘license’ Pch2 binding. In agreement with such a model, we found that during meiosis, Zip1 was required for the recruitment of Pch2 to *GAP1* (Figure 5E and Supplementary Figure 7A-C). As such, our results reveal an intricate connection between transcription, ORC/Orc1 function and meiosis-specific chromosome organization that allows the recruitment of a specific chromosomal pool of Pch2 (Figure 5F), an important regulator of meiotic chromosome metabolism.

## Discussion

The AAA+ protein Pch2 controls meiotic DSB formation, influences crossover recombination, mediates a meiotic G2/prophase checkpoint and is involved in chromosome reorganization upon chromosome synapsis (reviewed in [19]). For many functions, the chromosomal association of Pch2 has been postulated to be crucial. During meiotic G2/prophase Pch2 is enriched within the nucleolus/rDNA, and is also detected on synapsed chromosomes [17]. Chromosome synapsis is mediated by the dynamic polymerization of the synaptonemal complex (SC), whose formation is mostly nucleated at sites of crossover recombination [9-11]. Synapsis-dependent recruitment of Pch2 is abolished in cells that lack Zip1, the central element of the SC [12, 13, 17]. In addition to Zip1, other factors influence the chromosomal distribution of Pch2: Sir2 and Orc1 promote the nucleolar localization of Pch2 [21, 28], whereas Dot1 influences global chromosomal abundance of Pch2 [21, 22]. Here, we present a comprehensive analysis of the chromosomal association of Pch2 via genome-wide ChIP-seq. Surprisingly, we reveal that Pch2 is associated with a subset of actively transcribed RNAPII-dependent genes. We perform several experiments and analyses to ascertain that these binding patterns are biologically meaningful and not caused by ChIP-associated artefacts (see Supplementary results and discussion). Recruitment of Pch2 is dependent on active RNAPII-dependent transcription, but transcriptional strength *per se* is not a determining factor. For example, many actively transcribed genes do not recruit significant amounts of Pch2. Additional bioinformatics analysis failed to reveal any specific underlying characteristics of gene content driving Pch2 association. If not solely determined by transcriptional strength and specific DNA content, what are additional factors that influence Pch2 association? We find that, Orc1, a component of ORC, is required for Pch2 association with actively transcribed genes. The connection between Pch2 and Orc1 at genomic regions that are distinct from origins of replication further underscores the non-canonical role played by Orc1 during meiotic G2/prophase [47]. In addition, since Orc1 is also involved in nucleolar recruitment of Pch2 [28, 46], these findings hint at a common biochemical foundation that underlies recruitment of Pch2 to diverse chromatin environments. A recent study did not detect a role for Orc1 on the chromosomal (non-nucleolar) recruitment of Pch2 [46], and we speculate that differences in the experimental approaches that were used to interfere with Orc1 function might underlie this discrepancy. We show here that the Bromo-Adjacent Homology BAH-domain of Orc1, a nucleosome-binding module contributes to targeting of Pch2. BAH domains are readers of chromatin state [53]. A structural characteristic of a related BAH domain (*i.e.* that of budding yeast Sir3) is that nucleosome association is sensitive to Dot1-mediated Histone H3 K79 methylation (H3K79me) [56]. Dot1 activity (and H3-K79 methylation state) is important for Pch2 localization along chromosomes [22]. Dot1 activity is associated with active RNAPII transcriptional activity (reviewed in [60]), and RNAPII transcription-associated Dot1 activity may thus affect binding patterns of Pch2, potentially through Orc1 BAH domain-mediated nucleosome interactions. Of note, by analyzing correlations between our Pch2 data sets with genome wide maps of several histone modifications [61], we found a correlation between genome-wide Pch2 binding and H3K79-monomethylation patterns (data not shown). We thus speculate that epigenetic state (*i.e.* specific chromatin modifications) could contribute to Pch2 recruitment within euchromatin. Previous work has established that Pch2 localizes to the nucleolus [21], the site of RNAPI-driven rDNA transcription. The nucleolus is a membrane-less, self-organized nuclear compartment that exhibits properties of a liquid-liquid phase separated nuclear condensate (reviewed in [62]). Interestingly, recent work revealed the existence of RNAPII-specific nuclear condensates that influence transcriptional regulation (reviewed in [43]). In the future, it will be interesting to investigate whether the biochemical properties of transcriptional condensates relate to (shared) characteristics for the recruitment and function of Pch2 at RNAPI- and RNAPII-transcriptional hubs.

Meiotic chromosomes are organized into a typical loop-axis structure [3] [4]. Our genome-wide mapping revealed that the Pch2 binding patterns are strikingly distinct from the stereotypical binding patterns of meiotic axis factors (*i.e.* Hop1, Red1 and Rec8) [7, 8]. We suggest that the transcription-associated pool of Pch2 is not directly associated with chromosome axis sites and is instead associated with genes in loop regions. This is surprising, since *i*) the only identified client of Pch2, Hop1 is a component of the axis, and *ii*) assembly of the SC, a structure that is assembled directly on the axis factors, is important to allow Pch2 recruitment to chromosomes. The observed Pch2 binding pattern may indicate that the pool of Pch2 identified here plays a role that is distinct from the canonical role of Pch2 in the removal of the bulk of Hop1 from meiotic axis sites. Accordingly, interference with recruitment of this Pch2 population (via transcriptional inhibition) did not affect the chromosomal association of Hop1. We cannot rule out that incomplete inhibition of Pch2 association under the used conditions precludes us from exposing a contribution for this population of Pch2 in regulating Hop1. Nonetheless, also when considering the localization patterns of Pch2 described here, we speculate that more than one chromosomal populations of Pch2 might exist, of which one is recruited to transcriptionally active regions. In that scenario, a population of Pch2 that is directed to chromosome axis sites (and can conceivably not be detected using ChIP-based approaches) might be responsible for the majority of Hop1 removal upon chromosome synapsis.

Why and how does synapsis (*i.e.* Zip1 polymerization) contribute to the recruitment of the transcription-associated Pch2 population (Figure 5), and what is the role of the transcription-associated pool of Pch2? SC polymerization along synapsing chromosomes has been proposed to trigger mechanical reorganization [63], which might influence large-scale topological organization of loop-axis structures, and Pch2 localization. Indeed, previous work has identified a connection between topoisomerase II function, meiotic chromosome reorganization and Pch2 recruitment [23, 64]. It will be interesting to further understand the link between transcription, chromosome topology and organization and Pch2 recruitment. Many if not all of the roles Pch2 plays in meiosis have been attributed to its biochemical effects on Hop1 (reviewed in [19]). Although bulk Hop1 removal was not affected under conditions where a transcription-associated population of Pch2 was displaced (Figure 4), it remains possible that local Hop1 behavior is affected by this population. In addition, Pch2 impacts global DSB activity [38], and the specific recruitment of Pch2 to defined chromosomal regions might conceivably impact local DSB patterning. Finally, it will be interesting to consider whether the association of Pch2 with actively transcribed regions is in any way related to transcriptional regulation.

In conclusion, we have used genome-wide methodology to reveal a hitherto unknown relationship between Pch2, active transcription and Orc1, which influences the chromosome synapsis-driven recruitment of Pch2 to euchromatin during meiotic G2/prophase. Future work should increase our understanding of dynamic chromosome recruitment of this important regulator of meiotic DSB formation, recombination and checkpoint signaling.

## Acknowledgements

We thank members of the Vader and Bird laboratories for ideas and helpful discussions. We gratefully acknowledge financial support by the European Research Council (ERC Starting Grant URDNA, agreement nr. 638197, to GV), a CAPES-Humboldt fellowship from the Alexander von Humboldt Foundation (agreement nr. 99999.000021/2016-04, to RCS) and the Max Planck Society. We thank Andrea Musacchio (Max Planck Institute of Molecular Physiology, Dortmund, Germany) for ongoing support. We thank Jonna Heldrich and Andreas Hochwagen (NYU, New York City, USA) for sharing unpublished RNA-seq datasets. We acknowledge Stephen Bell (MIT, Cambridge, USA), Amy MacQueen (Wesleyan University, Middletown, USA) and Nancy Hollingsworth (Stony Brook University, Stony Brook, USA) for sharing reagents.

## Author contributions

Conception and experimental design: RCS and GV. Experimentation: RCS and MAVF. Computational analyses: RCS; Data analysis: GV and RCS; Supervision: GV; Manuscript: RCS and GV with input from MAVF.

