## Supplementary Data for "Active transcription and Orc1 drive chromatin association of the AAA+ ATPase Pch2 during meiotic G2/prophase"

**A chromatin-associated pool of the AAA+ ATPase Pch2 that is defined by active transcription and Orc1**

**Supplementary data**

**Supplementary results and discussion**

ChIP results can be plagued by artificial enrichments within regions of active transcription [1], and we performed several analyses and experiments to ascertain that the binding patterns we found for Pch2 are not a result of such artefactual effects. Based on these combined results we conclude that the Pch2 binding patterns observed here are a reflection of physiologically relevant effects.

1) We compared a “hyperChIPable” dataset obtained from yeast strains expressing GFP harboring a nuclear localization signal (NLS-GFP) [1] with our datasets. Peak calling was based on a highly stringent cut-off p-value of 1e-15, a significance used by others when reporting meiois-specific ChIP-seq data sets [2]. Re-analysis of the reported NLS-GFP data set [1] using the same cut-off value revealed no significant peaks. NLS-GFP crosslinks non-specifically with >273 peaks, which often are at genes encoding for ribosomal factors and tRNA genes (these account for ~70% of the peaks) [1]. Our identified list of Pch2 binding sites showed little overlap with the hyperChIPable dataset: only 19 (RNAPolII-transcribed) of a total of of 434 genes (*i.e.* ~4%) were reported within the NLS-GFP dataset (Supplementary Figure 2A). Accordingly, we detected no significant peaks at Pch2 at tRNAs genes ((Supplementary Figure 2B) and other non-coding RNAs, despite the fact that all the tRNA genes reported as artefactual ChIP signals are expressed during meiosis [3]. HyperChIPable regions show a positive correlation between ChIP signal and RNAPII activity [1]. In contrast, we detected a weak correlation between the peak shape score of individual Pch2-binding sites and expression level of the corresponding CDS (Pearson’s test, R^2^=0.0026, Supplementary Figure 1D). Inspection of the 19 binding sites that overlap between our dataset and the NLS-GFP dataset also revealed that 75% of these possess relatively moderate Pch2-enrichment scores in our ChIP-seq dataset.
2) HyperChIPpable regions have been suggested to originate from non-physiological crosslinking artefacts by which (*i.e.* an “inert” nuclear GFP shows crosslink-based enrichment) [4]. Based on such a scenario, any nuclear factor should exhibit significant artefactual binding. We used a strain expressing 3XFLAG-dCas9 (without the co-expression of a guide (g)RNA), an inert nuclear protein − containing an NLS − that is not expected to bind any specific loci without a gRNA) to query association with a peak identified in our Pch2 dataset, using ChIP-qPCR, and compared it to enrichment as seen with 3XFLAG-Pch2 (Supplementary Figure 2C and D). Although ChIP-qPCR readily detected a strong enrichment of 3XFLAG-Pch2 with this locus, 3XFLAG-dCas9 showed very minor enrichment, which was barely stronger than untagged controls.
3) We observed stronger peak scores in ChIP datasets from Pch2-E399Q as compared to wild type Pch2 (see above, and Supplementary Figure 1B). The same effect was also observed in ChIP-qPCR experiments (for example see Figure 1H). We also used a mutant of Pch2, lacking its NTD to investigate binding, and we observed a strong dependency of Pch2 ChIP signals on this domain (Figure 1H). All these experiments indicate that the association of Pch2 to chromatin behaves as would be expected, based on the known biochemical behavior of AAA+ enzymes [4]
4) We found that Pch2 binding depends on Orc1 function, and on the BAH domain of Orc1 (Figure 3)
5) We corroborated our ChIP-based analysis using independent cytological analyses (immunofluorescence on meiotic chromosome spreads). These analyses demonstrate that in cells where *i)* transcription was inhibited (Figure 2) or *ii)* Orc1 function was impaired (via usage of *orc1-161*) Pch2 recruitment was impaired (Figure 3).
6) Ectopic expression of Pch2 in mitosis does not lead to recruitment of Pch2 to sites of active transcription (Figure 5). Thus, under conditions where the interrogated regions were transcriptionally active (and Orc1/ORC was present), Pch2 was not recruited to chromosomes.
7) Finally, we showed that Pch2 recruitment depends on Zip1 function (Figure 5), as judged by ChIP-based analysis. This is in agreement with earlier cytological studies that have established a role for Zip1 in driving the chromosome recruitment of Pch2 [5].

**Supplementary materials and methods**

**ChIP-qPCR primers**

Primer efficiencies (PE) were calculated using standard procedures.

pGV2390 5’-AGAACGTCATCTCCGGAATCT-3’ PE=1.998

pGV2391 5’-TGGGCACGATGAGAGAAAGT-3’

pGV2577 5’-AAGCTTTTCATCCCAGCAGA- 3’ PE=1.991

pGV2578 5’-TTTTTGTCGTTGTTCGATTCA- 3’

pGV2583 5’-ACCATCAGGACTGGAAGTGG- 3’ PE=2.006

pGV2584 5’-CCTGTGGTGACGAAGAATCA- 3’

pGV2587 5’-CTTGAACAGCAGCACCGTAA- 3’ PE=1.992

pGV2588 5’-TGGACCCAGTTGAAAAGGTC- 3’

pGV2593 5’-TGGTGGTACAAGAAGCGTTG- 3’ PE=1.990

pGV2594 5’-ACATCGCCATTGACTCCACT- 3’

pGV2595 5’-TCGAAGACCGAACAGGAACT- 3’ PE=1.953

pGV2596 5’-ACAATTGGTCCAAAGCATCC- 3’

pGV2597 5’-GCGGACAAGAATGTGTCTGA- 3’ PE=2.001

pGV2598 5’-TCTGCTGTAAATGGCAAACG- 3’

pGV2599 5’-GAGGACGAGCTGGTGGAATA- 3’ PE=1.962

pGV2600 5’-TTCTGTTTCAGGTCCCCAAG- 3’

pGV2605 5’-GGGGTTGTTCCCTGATGATA- 3’ PE=1.927

pGV2606 5’-CTCCCTTCTCCAACCAATCC- 3’

pGV2717 5’-CAAGAAATGCAAACCGCTGC-3’ PE=2.019

pGV2718 5’-GGTCAATACCGGCAGATTCC-3’

pGV2747	5’-GCCTTAGTAACGGCGAGTGA-3’ PE=1.976

pGV2748 5’-CACGGGATTCTCACCCTCTA-3’

**Supplementary figure legends**

**Supplementary figure 1.**

A. Overview of expressed genes and binding sites for 3XFLAG-Pch2 and 3XFLAG-Pch2-E399Q during meiotic G2/prophase, as determined by ChIP-seq. Gene expression analysis was based on an mRNA dataset from [6]. B. Average peak shape scores for 3XFLAG-Pch2 and 3XFLAG-Pch2-E399Q during meiotic G2/prophase, as determined by ChIP-seq. C. Comparison between expression strength of Pch2-associated genes and the transcribed genes from the mRNA dataset [6], binned into high, medium and low expression strength (following previously established procedures [7]). D. Correlation between ChIP peak shape score of 3XFLAG-Pch2 individual binding sites and mRNA expression values [6]. E. ChIP-qPCR analysis of 3XFLAG-Pch2-E399Q at *PPR1* (yGV2390/yGV2391) and *GAP1* (yGV2597/yGV2598) during meiotic G2/prophase (4 hours). F. ChIP-qPCR analysis of active transcription (α-phosphoserine 5 Rpo21) at *PPR1* (yGV2390/yGV2391) and *GAP1* (yGV2597/yGV2598) during meiotic G2/prophase (4 hours).

**Supplementary figure 2.** A. Venn diagram comparing 3XFLAG-Pch2 binding peaks and HyperChIPpable regions as described by [1]. B. Whole genome average plotting of 3XFLAG-Pch2 and 3XFLAG-Pch2-E399Q binding peaks (log2). Datasets were aligned relative to the center of tRNAs. C. ChIP-qPCR analysis of 3XFLAG-Pch2 and 3XFLAG-dCas9 at *PPR1* (yGV2390/yGV2391) and *GAP1* (yGV2597/yGV2598) during meiotic G2/prophase (4 hours). Error bars represent standard error of at least three independent experiments performed in triplicate. D. Western blot analysis of 3XFLAG-Pch2 and 3XFLAG-dCas9 as used in C.

**Supplementary figure 3.** A. mRNA quantification of *GAP1* (yGV2597/yGV2598)*/18S (*yGV2717/yGV2718) in cells (wild type or 3XFLAG-Pch2-E399Q) treated with 20% EtOH or 1,10- Phenanthroline. B. Schematic of 1,10- Phenanthroline treatment regimen as used for C-D. C. Western blot analysis of 3XFLAG-Pch2 in cells (wild type or 3XFLAG-Pch2-E399Q) treated with 20% EtOH or 1,10- Phenanthroline. D. ChIP-qPCR analysis of 3XFLAG-Pch2-E399Q at *PPR1* (yGV2390/yGV2391)*, GAP1* (yGV2597/yGV2598) in cells (wild type or 3XFLAG-Pch2-E399Q) treated with 20% EtOH or 1,10- Phenanthroline. Error bars represent standard error of at least three independent experiments performed in triplicate. E. Western blot analysis of Rpo21 (α-FRB or α-Rpo21) in wild type and *rpo21-FRB* anchor away strains, upon treatment with DMSO or rapamycin, treated as described in Figure 2C.

**Supplementary figure 4.** A. ChIP-qPCR analysis of ORC (α-ORC) along the *GAP1* locus during meiotic G2/prophase (4 hours). Primers pairs 1: yGV2595/yGV2596, 2: yGV2597/yGV2598, 3: yGV2599/yGV2600. Error bars represent standard error of at least three independent experiments performed in triplicate. B. Western blot analysis of Orc1 and Orc2 (α-ORC) in *ORC1* and *orc1-161* cells. Experiment was performed at 23°C. C. Flow cytometric analysis of *ORC1* and *orc1-161* cells (wild type, or expressing or 3XHA-Pch2-E399Q). Time (hours) after induction into the meiotic program is indicated. Experiment was performed at 23°C. D. ChIP-qPCR analysis of Orc2-TAP at *PPR1* (yGV2390/yGV2391)*, GAP1* (yGV2597/yGV2598) and *ARS1116* (yGV2577/yGV2578) in *ORC1* and *orc1-161* cells. Experiment was performed at 23°C. Error bars represent standard error of at least three independent experiments performed in triplicate. E. Western blot analysis of 3XHA-Pch2 (α-HA) in *ORC1* and *orc1-161* cells. Hours after induction into the meiotic program indicated. Experiment was performed at 23°C. F. Western blot analysis of Orc1 and Orc2 (α-ORC) in *ORC2* and *orc2-1* cells. Experiment was performed at 30°C. G. Flow cytometric analysis of *ORC2* and *orc2-1* cells (wild type, or expressing or 3XHA-Pch2-E399Q). Hours after induction into the meiotic program indicated. Experiment is performed at 30°C. H. ChIP-qPCR analysis of 3XHA-Pch2-E399Q at *PPR1* (yGV2390/yGV2391) and *GAP1* (yGV2597/yGV2598) in *ORC2* and *orc2-1* cells. Experiment was performed at 30°C. Error bars represent standard error of at least three independent experiments performed in triplicate. I. Western blot analysis of 3XHA-Pch2 and Orc1 (α-HA and α-ORC) in *ORC1* and *orc1Δbah* cells. Hours after induction into the meiotic program indicated. J. Flow cytometric analysis of *ORC1* and *orc1Δbah* cells (wild type, or expressing 3XHA-Pch2-E399Q). Hours after induction into the meiotic program indicated.

**Supplementary figure 5.** A and B. Representative images of ChIP-seq binding patterns for 3XFLAG-Pch2, Hop1, Red1 and Rec8. Data for Hop1, Red1 and Rec8 are from [2]. Shown is entire chromosome *I* (A) and a region of chromosome *III (B)* (chromosomal coordinates (kb) are indicated).

**Supplementary figure 6.** A. Schematic of treatment regimen used for anchor away experiments as used for B-E. B. Immunofluorescence of meiotic chromosome spreads in the 3XHA-Pch2 expressing *rpo21-FRB* anchor away cells, treated with DMSO or rapamycin as in A. Chromosome synapsis was assessed by α-Gmc2 staining. C. Quantification of B. **** indicates a significance of p≤ 0.0001, Mann-Whitney U test. D. Hop1 immunofluorescence of meiotic chromosome spreads in *rpo21-FRB* anchor away cells, treated with DMSO or rapamycin as in A. Chromosome synapsis was assessed by α-Gmc2 staining. E. Quantification of D. n.s. indicates p>0.05, Mann-Whitney U test. F. Representative images of Hop1 ChIP-seq binding patterns in wild type and pch2*Δ* from [8]. Chromosome coordinates and primer pairs are indicated. G. ChIP-qPCR analysis of Hop1 in *rpo21-FRB* anchor away strains, upon treatment with DMSO or rapamycin, treated as described in Figure 2C. Primers are described in [8]. Error bars represent standard error of at least three independent experiments performed in triplicate.

**Supplementary figure 7.** A. mRNA quantification of *GAP1*(yGV2597/yGV2598) *(*yGV2717/yGV2718) *or PPR1*(yGV2390/yGV2391)*/*β-Actin (*ACT1;* yGV2747/yGV2748) in wild type or *zip1Δ* cells during meiotic G2/prophase (4 hours). B. Flow cytometric analysis of wild type and *zip1Δ* cells (wild type, or expressing 3XFLAG-Pch2-E399Q). Time (hours) after induction into the meiotic program is indicated. C. Western blot analysis of 3XFLAG-Pch2 and Zip1 (α-FLAG and α-Zip1) in wild type and *zip1Δ* cells. Time (hours) after induction into the meiotic program is indicated.

**Yeast strains**

All strains, except yGV104 and yGV2941 which are of the W303 background, are derived from the SK1 background.

yGV**49**	*MATa/MATalpha, ho::LYS2, lys2, ura3, leu2::hisG, his4B::LEU2, ARG4/*

*arg4-Bgl II*

yGV104	*MATa, ade2-1, leu2-3, ura3, trp1-1, his3-11,15, can1-100, GAL, psi+*

yGV933	*MATa/MATα, ho::LYS2, lys2, ura3, leu2::hisG, trp1::hisG, his3::hisG,	his4B::LEU2, arg4-Bgl II, pch2::URA3:pPCH2(300bp):3HA-PCH2*

yGV1185   	MATa/ MATalpha, ho::LYS2, lys2, ura3, leu2::hisG, TRP1, HIS3,  arg4-Bgl	II, pch2::URA3:pPCH2(300bp):3HA-PCH2, orc1::orc1-161 (ts-allele)

yGV**1465**	*MATa/MATalpha,  ho::LYS2, lys2, ura3, leu2::hisG, his3::hisG, trp1::hisG*

*orc1::TRP1, ura3::orc1ΔNTD(1-235)::URA3*

yGV1506	*MATa/MATα, ho::LYS2, lys2, ura3, leu2::hisG, TRP1, HIS3,*

*his4B::LEU2, pch2::URA3:pPCH2(300bp):3HA-PCH2, orc1::ORC1-TAP::HIS3*

yGV1508	*MATa/MATα, ho::LYS2, lys2, ura3, leu2::hisG, HIS3, trp1::hisG, his4B::LEU2,	orc2::ORC2-TAP::HIS3, pch2::URA3:pPCH2(300bp):3HA-PCH2*

yGV1945	*MATa/MATα, ho::LYS2, lys2, ura3, leu2::hisG, HIS3, trp1::hisG, his4B::LEU2,	orc2::ORC2-TAP::HIS3, pch2::URA3:pPCH2(300bp):3HA-PCH2, orc1::orc1-161*

yGV**2086**	*MATa/MATalpha, ho::LYS2, lys2, ura3, leu2::hisG, TRP, HIS3*

*his4B::LEU2, arg4-Bgl II, pch2::URA3:pPCH2(300bp):3HA-PCH2-E399Q*

yGV2099	*MATa/MATα, ho::LYS2, lys2, ura3, leu2::hisG, trp1::hisG, his3::hisG, arg4-	Nsp/arg4, his4X::LEU2-(BGV)-URA3, pch2Δ::KanMX, orc1::ORC1-TAP::HIS3*

yGV2234	*MATa/MATalpha, ho::LYS2, lys2, ura3, leu2::hisG, TRP1, HIS3*

*his4X, ARG4/ arg4-BglII, pch2::URA3:pPCH2(300bp):3HA-	PCH2,  ndt80Δ::TRP1*

yGV2447	*MATa/MATα, ho::LYS2, lys2, ura3, leu2::hisG, his3::hisG, trp1::hisG,	his4X::LEU2-	URA3, trp1::pPch2::TRP1, ARG4, pch2Δ::KanMX, ndt80Δ::LEU2*

yGV2249	*MATa/MATα, ho:LYS2, lys2, ura3, leu2::hisG, TRP1, ARG4 ndt80Δ::TRP1,	pch2::URA3:pPCH2(300bp):3HA-pch2-E399Q*

yGV2875	*MATa/MATα, ho::LYS2, lys2, leu2::hisG, his3::hisG, ura3, trp1:pPCH2-FLAG-	6GLY-ΔNTD (1-242)-Pch2::TRP1, pch2Δ::KanMX, ARG4, ndt80Δ::LEU2*

yGV2889	*MATa/MATα, ho::LYS2, lys2, leu2::hisG, his4X::LEU2-URA3, his3::hisG, ura3,	trp1:pPCH2-3FLAG-6GLY-PCH2::TRP1, pch2Δ::KanMX, ARG4, ndt80Δ::TRP1*

yGV2899	*MATa/MATα, ho::LYS2, lys2, leu2::hisG, his3::hisG, ura3, trp1:pPCH2-3FLAG-	6GLY-PCH2::TRP1, pch2Δ::KanMX, ARG4, orc1::TRP1, ura3::orc1ΔNTD(1-	235)-TAP::HIS3::URA3*

yGV2919	*MATa/MATα, ho::LYS2, lys2, leu2::hisG, his4X::LEU2-URA3, HIS3 , ura3,	trp1::hisG, pch2Δ::KanMX, ARG4, trp1:pPch2-FLAG-6XGLY-pch2	E399Q::TRP1, ndt80Δ::TRP1*

yGV2941	*MATa, ade2-1, leu2-3, ura3, trp1-1, his3-11,15, can1-100, GAL, psi+,	ura3:pGAL10-3HA-pch2-E399Q::URA3*

yGV2950	*MATa/MATalpha, ho::LYS2, lys2, ura3, leu2::hisG, his3::hisG,	trp1::hisG, ARG4, trp::p11_pHOP1_3xFlag-dCas9::TRP1*

yGV3304	*MATa/MATα, ho::LYS2, lys2, leu2::hisG, his4X::LEU2-URA3, HIS3, ARG4,	ura3, trp1::hisG, pch2Δ::KanMX, ARG4, trp1:pPch2-FLAG-6XGLY-pch2	E399Q::TRP1, ndt80Δ::TRP1, orc1::orc1-161*

YGV3320	*MATa/MATα, ho::LYS2, lys2, ura3, leu2::hisG, his3::hisG, trp1::hisG,	his4X::LEU2,	trp1::pPch2::TRP1, ARG4, pch2Δ::KanMX, ndt80Δ::LEU2,	orc1::orc1-161*

yGV3384	*MATa/MATα, ho::LYS2, lys2, ura3, leu2::hisG, TRP, arg4-Bgl II*

*pch2::URA3:pPCH2(300bp):3HA-pch2-E399Q, orc1::TRP1, ura3::ORC1	TAP::HIS::URA3*

yGV3422	*MATa/MATα, ho::LYS2, lys2, leu2::hisG, his4X::LEU2-URA3, HIS3, ura3,	trp1::hisG, pch2Δ::KanMX, ARG4, trp1:pPch2-3XFLAG-6XGLY-pch2	E399Q::TRP1, ndt80Δ::TRP1, zip1Δ::NatMX4*

yGV3598	*MATa/MATα, ho::LYS2, lys2, leu2::hisG, his4X::LEU2-URA3, HIS3, ura3,	trp1::hisG, pch2Δ::KanMX, ARG4, trp1:pPch2-3XFLAG-6XGLY-pch2	E399Q::TRP1, ndt80Δ::TRP1, dot1Δ::NatMX4, zip1Δ::NatMX4*

yGV3898	*MATa/MATα, ho::LYS2, lys2, ura3, leu2::hisG, his3::hisG, trp1::hisG, , RPL13A-	2XFKBP12::TRP1, fpr1::KanMX4, tor1-1::HIS3, ndt80Δ::NatMX4,	pch2::URA3:pPCH2(300bp):3HA-PCH2*

yGV3943	*MATa/MATα, ho::LYS2, lys2, ura3, leu2::hisG, his3::hisG, trp1::hisG, RPO21-	FRB::KANMX6, RPL13A-2XFKBP12::TRP1, fpr1::KanMX4, tor1-1::HIS3,	ndt80Δ::NatMX4, pch2::URA3:pPCH2(300bp):3HA-PCH2*

yGV4034	*MATa/MATα, ho::LYS2, lys2, ura3, his4B::LEU2, ARG4,	pch2::URA3:pPCH2(300bp):3HA-PCH2-E399Q, orc2-1, ndt80Δ::NatMX4*

yGV4041	*MATa/MATα, ho::LYS2, lys2, ura3, TRP1, ARG4, orc2-1, ndt80Δ::NatMX4*

**yGV4161**	*MATa/ MATalpha, ho::LYS2, lys2, ura3, leu2::hisG, TRP1, ARG4,	pch2::URA3:pPCH2(300bp):3HA-pch2-E399Q****,*** *orc1::TRP1,	ura3::orc1ΔNTD(1-235)::URA3*

**Yeast strains used per figure:**

1B:	yGV2447, yGV2889 and yGV2919

1C:	yGV2889 and yGV2919

1D:	yGV2889

1E:	yGV2889 and yGV2919

1F:	yGV2447 and yGV2919

1G and H:	yGV2447, yGV2875, yGV2889 and yGV2919

2B:	yGV3943 and yGV3898

2D and E:	yGV3943

2F and G:	yGV3898 and yGV3943

2H and I:	GV3943

3B:	yGV2889 and yGV2919

3C:	yGV2447 and yGV2919

3D:	yGV1506, yGV1508 and yGV2447

3E:	yGV2447 and yGV2919, yGV3304 and yGV3320

3F and G:	yGV933 and yGV1185

3H:	yGV49, yGV1465, yGV2086 and yGV4116

4A:	yGV2889

4C and D:	yGV3943

4F:	yGV3943

5B:	yGV2941, yGV2086

5C and D:	yGV49, yGV104, yGV2941, yGV2086

5E:	yGV2447, yGV2919, yGV3598

**Yeast strains used per supplementary figure:**

S1B:	yGV2889 and yGV2919

S1C and D:	yGV2889

S1E:	yGV2447

S1F:	yGV2447 and yGV2919

S2B:	yGV2889 and yGV2919

S2C and D:	yGV49, yGV2447, yGV2889, and yGV2950

S3A-C:	yGV2447 and yGV2919

S3D:	yGV2234, yGV3898 and yGV3943

S4A:	yGV2919

S4B:	yGV2919 and yGV3304

S4C:	yGV2447, yGV2919, yGV3304 and yGV3320

S4D	yGV933, yGV1185, yGV1508 and yGV1945

S4E:	yGV933 and yGV1185

S4F:	yGV49, yGV4034, yGV4041

S4G and H:	yGV49, yGV2086, yGV4034, yGV4041

S4I:	yGV2086, yGV4034, yGV4041

S4J:	yGV49, yGV1465, yGV2086 and yGV4161

S5A and B:	yGV2889

S6B-G:	yGV3943
