## Supplementary Table 1 for "Active transcription and Orc1 drive chromatin association of the AAA+ ATPase Pch2 during meiotic G2/prophase"

Lists of identified Pch2 regions, associated with CDS. A. 434 shared binding regions between 3XFLAG-Pch2 and 3XFLAG-Pch2-E399Q, B.91 Additional binding sites identified specifically for 3XFLAG-Pch2-E399Q. Chromosomal (SGD) coordinates are indicated.

A. 3X-FLAG-Pch2 and 3XFLAG-Pch2-E399Q

| **Chromosome** | **Gene name** | **Start (bp)** | **End(bp)** |
| --- | --- | --- | --- |
| 1 | *FLO9* | 30639 | 31053 |
| 1 | *GDH3* | 32044 | 32425 |
| 1 | *BDH2* | 33859 | 34259 |
| 1 | *BDH1* | 35490 | 35886 |
| 1 | *PEX22* | 43151 | 43536 |
| 1 | *ACS1* | 44271 | 44733 |
| 1 | *CDC19* | 139454 | 139842 |
| 1 | *FUN19* | 140137 | 140701 |
| 1 | *SSA1* | 141118 | 141505 |
| 1 | *EFB1* | 142585 | 143048 |
| 1 | *YAT1* | 190456 | 190846 |
| 2 | *ECM21* | 37026 | 37427 |
| 2 | *ATP1* | 37922 | 38456 |
| 2 | *SEF1* | 100524 | 101064 |
| 2 | *PSY4* | 135044 | 135498 |
| 2 | *ECM13* | 137025 | 137422 |
| 2 | *SHE1* | 163162 | 163679 |
| 2 | *ACH1* | 194637 | 195147 |
| 2 | *UTP20* | 235316 | 235730 |
| 2 | *HTB2* | 236585 | 237022 |
| 2 | *ECM15* | 238133 | 238544 |
| 2 | *FLR1* | 255256 | 255666 |
| 2 | *HHT1* | 256430 | 256829 |
| 2 | *CST26* | 322365 | 322728 |
| 2 | *TCM62* | 323263 | 323817 |
| 2 | *QDR3* | 324785 | 325176 |
| 2 | *AAC3* | 418731 | 419112 |
| 2 | *TEF2* | 478336 | 478817 |
| 2 | *YBR085C-A* | 637272 | 637651 |
| 2 | *DUR1,2* | 638491 | 639010 |
| 2 | *OM14* | 639484 | 639867 |
| 2 | *FTH1* | 640176 | 640594 |
| 2 | *DUR1,2* | 641520 | 641814 |
| 2 | *ROT2* | 679653 | 680030 |
| 2 | *YBR230W-A* | 680842 | 681406 |
| 2 | *YBR255C-A* | 727570 | 728006 |
| 2 | *RIB5* | 797069 | 797529 |
| 2 | *PCA1* | 797858 | 798329 |
| 3 | *MGR1* | 49454 | 49879 |
| 3 | *GLK1* | 51223 | 51632 |
| 3 | *GID7* | 51838 | 52333 |
| 3 | *RRP7* | 66588 | 66995 |
| 3 | *HIS4* | 67700 | 68263 |
| 3 | *FRM2* | 76483 | 76863 |
| 3 | *KCC4* | 77038 | 77600 |
| 3 | *AGP1* | 78767 | 79152 |
| 3 | *LEU2* | 91836 | 92388 |
| 3 | *SGF29* | 105055 | 105501 |
| 3 | *YCL001W-B* | 114277 | 114665 |
| 3 | *YCP4* | 121464 | 121984 |
| 3 | *RVS161* | 132477 | 132932 |
| 3 | *PGK1* | 138061 | 138600 |
| 3 | *POL4* | 138600 | 138926 |
| 3 | *SLM5* | 162605 | 163050 |
| 4 | *ADY3* | 29794 | 30319 |
| 4 | *YDL199C* | 104025 | 104466 |
| 4 | *LYS20* | 134214 | 134624 |
| 4 | *YCR024C-B* | 149683 | 150126 |
| 4 | *GUD1* | 151250 | 151563 |
| 4 | *GGC1* | 152143 | 152515 |
| 4 | *GLT1* | 153388 | 153914 |
| 4 | *PAR32* | 154378 | 154839 |
| 4 | *PPH21* | 221494 | 221876 |
| 4 | *RRP42* | 264503 | 264903 |
| 4 | *BDF2* | 331248 | 331631 |
| 4 | *COX9* | 332252 | 332806 |
| 4 | *CBS1* | 334026 | 334424 |
| 4 | *NAT1* | 382521 | 383080 |
| 4 | *GPD1* | 412398 | 412833 |
| 4 | *RMD1* | 449669 | 450066 |
| 4 | *MRH1* | 508668 | 509041 |
| 4 | *KRS1* | 527736 | 528121 |
| 4 | *ENA2* | 538178 | 538566 |
| 4 | *VMS1* | 555908 | 556313 |
| 4 | *TPS2* | 595711 | 596222 |
| 4 | *SED1* | 601322 | 601705 |
| 4 | *YDR089W* | 625331 | 625810 |
| 4 | *YDR098C-A* | 652923 | 653297 |
| 4 | *BMH2* | 653996 | 654543 |
| 4 | *VBA4* | 690982 | 691401 |
| 4 | *YDR132C* | 721022 | 721436 |
| 4 | *ENT5* | 768521 | 768927 |
| 4 | *HOM2* | 770719 | 771235 |
| 4 | *CRF1* | 914321 | 914703 |
| 4 | *HTA1* | 915607 | 916011 |
| 4 | *RKM4* | 972301 | 972832 |
| 4 | *HSP78* | 973270 | 973701 |
| 4 | *BSC2* | 1013114 | 1013616 |
| 4 | *YRA1* | 1236717 | 1237265 |
| 4 | *YDR381C-A* | 1237526 | 1238013 |
| 4 | *NKP1* | 1241456 | 1241835 |
| 4 | *NPL3* | 1328919 | 1329456 |
| 4 | *GPI17* | 1329457 | 1330021 |
| 4 | *DIG2* | 1418909 | 1419374 |
| 4 | *PHO8* | 1419587 | 1419993 |
| 4 | *PLM2* | 1453692 | 1454255 |
| 4 | *AGE1* | 1489392 | 1489796 |
| 4 | *HLR1* | 1495985 | 1496528 |
| 4 | *HSP31* | 1503627 | 1504011 |
| 4 | *FIT1* | 1504300 | 1504851 |
| 5 | *DLD3* | 16650 | 17091 |
| 5 | *SIT1* | 27938 | 28488 |
| 5 | *CAN1* | 28692 | 29178 |
| 5 | *AVT2* | 32038 | 32449 |
| 5 | *PRB1* | 32764 | 33281 |
| 5 | *CIN8* | 40534 | 40955 |
| 5 | *GLY1* | 41308 | 41828 |
| 5 | *FRD1* | 67558 | 67988 |
| 5 | *MTC7* | 68199 | 68759 |
| 5 | *HYP2* | 85800 | 86183 |
| 5 | *GCD11* | 139184 | 139559 |
| 5 | *YAT2* | 202524 | 203035 |
| 5 | *GCN4* | 203775 | 204136 |
| 5 | *EDC2* | 222769 | 223157 |
| 5 | *SAH1* | 237644 | 238040 |
| 5 | *JHD1* | 256549 | 257053 |
| 5 | *HOM3* | 257332 | 257742 |
| 5 | *PIC2* | 260638 | 261022 |
| 5 | *GIP2* | 265157 | 265600 |
| 5 | *HIS1* | 267393 | 267831 |
| 5 | *ICL1* | 285526 | 286081 |
| 5 | *VHR2* | 286081 | 286500 |
| 5 | *MOT2* | 294386 | 294778 |
| 5 | *ARG5,6* | 295583 | 296145 |
| 5 | *RNR1* | 296390 | 296867 |
| 5 | *VTC1* | 302877 | 303269 |
| 5 | *ALD5* | 304277 | 304744 |
| 5 | *RPS24A* | 304882 | 305442 |
| 5 | *SRG1* | 322346 | 322768 |
| 5 | *TRP2* | 340191 | 340585 |
| 5 | *MET6* | 341093 | 341602 |
| 5 | *UBP5* | 460989 | 461548 |
| 5 | *DMC1* | 549025 | 549521 |
| 6 | *YFL040W* | 53998 | 54482 |
| 6 | *HAC1* | 75309 | 75837 |
| 6 | *GAT1* | 96228 | 96647 |
| 6 | *PAU5* | 96948 | 97326 |
| 6 | *YFR032C-B* | 224380 | 224762 |
| 6 | *YFR057W* | 264525 | 264951 |
| 7 | *ZRT1* | 21480 | 21958 |
| 7 | *ARO8* | 116651 | 117101 |
| 7 | *STR3* | 156696 | 157115 |
| 7 | *SCS3* | 272497 | 273048 |
| 7 | *RPL28* | 311279 | 311702 |
| 7 | *MMS2* | 346025 | 346463 |
| 7 | *DBP3* | 363201 | 363640 |
| 7 | *PYC1* | 385621 | 386080 |
| 7 | *DUO1* | 386660 | 387177 |
| 7 | *HNM1* | 388220 | 388605 |
| 7 | *OLE1* | 398917 | 399478 |
| 7 | *MPO1* | 477920 | 478424 |
| 7 | *LEU1* | 480610 | 481014 |
| 7 | *PMA1* | 482086 | 482520 |
| 7 | *UGA1* | 525403 | 525925 |
| 7 | *VMA7* | 526117 | 526496 |
| 7 | *ERG25* | 610957 | 611367 |
| 7 | *ADE6* | 623002 | 623348 |
| 7 | *YGR066C* | 624220 | 624755 |
| 7 | *RPL11B* | 649968 | 650407 |
| 7 | *MEP1* | 731788 | 732155 |
| 7 | *PIL1* | 732290 | 732735 |
| 7 | *ASN2* | 741168 | 741580 |
| 7 | *MEP1* | 742970 | 743353 |
| 7 | *YGR125W* | 744742 | 745235 |
| 7 | *NSR1* | 808686 | 809091 |
| 7 | *RTS3* | 847113 | 847454 |
| 7 | *YGR174W-A* | 847671 | 848130 |
| 7 | *TIM13* | 858992 | 859399 |
| 7 | *TDH3* | 882961 | 883525 |
| 7 | *TOS2* | 939306 | 939702 |
| 7 | *KEL2* | 969329 | 969741 |
| 7 | *MPC3* | 978216 | 978620 |
| 7 | *LSC2* | 1000991 | 1001440 |
| 7 | *ENO1* | 1001702 | 1002112 |
| 8 | *SBP1* | 35370 | 35781 |
| 8 | *NPR3* | 56754 | 57225 |
| 8 | *RIM4* | 57939 | 58323 |
| 8 | *DUR3* | 72285 | 72656 |
| 8 | *ERG11* | 73030 | 73419 |
| 8 | *YHL017W* | 73586 | 74103 |
| 8 | *STP2* | 120085 | 120559 |
| 8 | *SOD2* | 120783 | 121285 |
| 8 | *YHR007C-A* | 123029 | 123420 |
| 8 | *YSC83* | 140367 | 140838 |
| 8 | *PCL5* | 237231 | 237618 |
| 8 | *HXT1* | 294713 | 295237 |
| 8 | *YCK1* | 374566 | 374989 |
| 8 | *MPC2* | 423073 | 423485 |
| 8 | *ENO2* | 451711 | 452099 |
| 8 | *BAT1* | 518053 | 518611 |
| 9 | *YIL169C* | 23301 | 23734 |
| 9 | *HXT12* | 24693 | 25098 |
| 9 | *OM45* | 93967 | 94347 |
| 9 | *SPO22* | 222937 | 225954 |
| 9 | *HOP1* | 226602 | 228419 |
| 9 | *GPP1* | 255274 | 255750 |
| 9 | *ULP2* | 292801 | 293175 |
| 9 | *SSM4* | 293389 | 293850 |
| 9 | *RPL34B* | 294931 | 295452 |
| 9 | *YIR018C-A* | 390218 | 390612 |
| 9 | *FLO11* | 391801 | 392194 |
| 9 | *YIR018C-A* | 393084 | 393538 |
| 9 | *SSM4* | 408716 | 409261 |
| 9 | *DAL4* | 409309 | 409806 |
| 9 | *DAL2* | 411265 | 411703 |
| 9 | *DCG1* | 413245 | 413680 |
| 9 | *DAL3* | 413966 | 414497 |
| 9 | *DAL7* | 415060 | 415436 |
| 9 | *MGA2* | 419882 | 420445 |
| 9 | *PAU15* | 435468 | 435873 |
| 10 | *RPS14B* | 73929 | 74474 |
| 10 | *YJL171C* | 97943 | 98508 |
| 10 | *CPS1* | 98610 | 99013 |
| 10 | *QCR8* | 106513 | 106899 |
| 10 | *HSP150* | 121043 | 121497 |
| 10 | *VPS35* | 135263 | 135651 |
| 10 | *INO1* | 136339 | 136747 |
| 10 | *LCB3* | 159457 | 159861 |
| 10 | *URA2* | 172624 | 173026 |
| 10 | *SET4* | 221760 | 222263 |
| 10 | *IME2* | 222613 | 223092 |
| 10 | *PAM16* | 228029 | 228418 |
| 10 | *PRY1* | 265974 | 266456 |
| 10 | *ARG3* | 266456 | 266869 |
| 10 | *SIP4* | 267779 | 268232 |
| 10 | *ARG3* | 269231 | 269616 |
| 10 | *SCP160* | 290190 | 290585 |
| 10 | *PRE3* | 436043 | 436422 |
| 10 | *VPS55* | 519716 | 520220 |
| 10 | *SSC1* | 521083 | 521468 |
| 10 | *CDC11* | 577560 | 578124 |
| 10 | *FIP1* | 604752 | 605159 |
| 10 | *MIR1* | 605252 | 605667 |
| 10 | *URA8* | 622536 | 622961 |
| 10 | *IME1* | 629668 | 630217 |
| 10 | *CPA2* | 630740 | 631198 |
| 10 | *ABM1* | 631481 | 631856 |
| 10 | *SOD1* | 632234 | 632690 |
| 10 | *YJR115W* | 640238 | 640619 |
| 10 | *ATP2* | 647786 | 648034 |
| 10 | *IBA57* | 648451 | 648838 |
| 10 | *DAN1* | 713843 | 714216 |
| 10 | *DAL5* | 720595 | 720942 |
| 11 | *SRY1* | 21500 | 21906 |
| 11 | *JEN1* | 22621 | 23006 |
| 11 | *JEN1* | 23335 | 23827 |
| 11 | *DAN4* | 100718 | 101208 |
| 11 | *JEN1* | 102909 | 103318 |
| 11 | *URA1* | 104469 | 104848 |
| 11 | *PRS1* | 105265 | 105742 |
| 11 | *FAS1* | 105742 | 106264 |
| 11 | *DBR1* | 170491 | 171052 |
| 11 | *HAP4* | 232755 | 233112 |
| 11 | *MDH1* | 279528 | 279947 |
| 11 | *BLI1* | 326689 | 327235 |
| 11 | *PHD1* | 356808 | 357340 |
| 11 | *DAL80* | 507168 | 507591 |
| 11 | *GAP1* | 516105 | 516506 |
| 11 | *UTH1* | 519684 | 520075 |
| 11 | *YKR045C* | 524448 | 524896 |
| 11 | *PCK1* | 631782 | 632217 |
| 12 | *ATG10* | 53278 | 53762 |
| 12 | *VPS13* | 64159 | 64636 |
| 12 | *UBI4* | 64707 | 65135 |
| 12 | *FRA1* | 81791 | 82195 |
| 12 | *ISA1* | 87574 | 88041 |
| 12 | *PAU17* | 96934 | 97327 |
| 12 | *SPA2* | 106694 | 107256 |
| 12 | *KNS1* | 108474 | 108840 |
| 12 | *SED5* | 197223 | 197701 |
| 12 | *ADE16* | 198920 | 199379 |
| 12 | *TRX1* | 232981 | 233489 |
| 12 | *ERG3* | 254200 | 254663 |
| 12 | *MNL2* | 258579 | 258964 |
| 12 | *RPL10* | 283079 | 283503 |
| 12 | *AHP1* | 368887 | 369398 |
| 12 | *CCW12* | 369710 | 370088 |
| 12 | *SLX4* | 415930 | 416311 |
| 12 | *TIS11* | 425317 | 425788 |
| 12 | *PUT1* | 426045 | 426602 |
| 12 | *ACS2* | 445895 | 446330 |
| 12 | *YLR152C* | 446774 | 447334 |
| 12 | *RPS31* | 499563 | 499966 |
| 12 | *IDP2* | 505250 | 505691 |
| 12 | *SAM1* | 515461 | 515913 |
| 12 | *QRI5* | 552969 | 553355 |
| 12 | *HMX1* | 568755 | 569266 |
| 12 | *CDC123* | 569307 | 569721 |
| 12 | *FRE1* | 570164 | 570554 |
| 12 | *THI7* | 613184 | 613585 |
| 12 | *FAR10* | 637172 | 637555 |
| 12 | *SSP120* | 637839 | 638395 |
| 12 | *YEF3* | 639328 | 639879 |
| 12 | *YLR257W* | 659218 | 659666 |
| 12 | *ACO1* | 736728 | 737203 |
| 12 | *CDA1* | 746715 | 747094 |
| 12 | *YLR326W* | 783014 | 783415 |
| 12 | *RPP0* | 806115 | 806529 |
| 12 | *TAL1* | 838583 | 839147 |
| 12 | *CCW14* | 903909 | 904400 |
| 12 | *CTR3* | 947456 | 947990 |
| 12 | *CAR2* | 1012891 | 1013455 |
| 13 | *RSC9* | 19699 | 20161 |
| 13 | *TUB3* | 24921 | 25321 |
| 13 | *ERG13* | 87289 | 87781 |
| 13 | *PHO84* | 88503 | 89066 |
| 13 | *RPM2* | 89421 | 89864 |
| 13 | *PRE8* | 90202 | 90641 |
| 13 | *RPM2* | 91001 | 91511 |
| 13 | *WAR1* | 116070 | 116570 |
| 13 | *SML1* | 159418 | 159807 |
| 13 | *CMP2* | 192975 | 193397 |
| 13 | *VPS71* | 193630 | 194029 |
| 13 | *TSA1* | 220291 | 220668 |
| 13 | *YPT7* | 267940 | 268323 |
| 13 | *MIX17* | 272301 | 272698 |
| 13 | *HXT2* | 288431 | 288858 |
| 13 | *snR78* | 297361 | 297904 |
| 13 | *BUD22* | 301169 | 301494 |
| 13 | *ERG5* | 301746 | 302297 |
| 13 | *FET3* | 389208 | 389764 |
| 13 | *NAM7* | 430284 | 430688 |
| 13 | *YMR087W* | 444084 | 444289 |
| 13 | *VBA1* | 444461 | 444889 |
| 13 | *YPK2* | 476317 | 476757 |
| 13 | *ILV2* | 484262 | 484692 |
| 13 | *MYO5* | 485628 | 486111 |
| 13 | *YMR144W* | 554749 | 555233 |
| 13 | *NDE1* | 555556 | 555937 |
| 13 | *YMR182W-A* | 626298 | 626690 |
| 13 | *HSC82* | 632479 | 633038 |
| 13 | *YMR187C* | 633038 | 633475 |
| 13 | *SGS1* | 633761 | 634039 |
| 13 | *YMR196W* | 637857 | 638229 |
| 13 | *GCV2* | 639466 | 639846 |
| 13 | *ICY1* | 654175 | 654587 |
| 13 | *ERG2* | 667639 | 668129 |
| 13 | *INP1* | 673986 | 674380 |
| 13 | *YMR196W* | 760119 | 760573 |
| 13 | *FAA4* | 761361 | 761902 |
| 13 | *HOR7* | 774778 | 775196 |
| 13 | *YME2* | 873447 | 873827 |
| 13 | *ADH6* | 913106 | 913488 |
| 13 | *FET4* | 914033 | 914590 |
| 14 | *MRPL17* | 174197 | 174584 |
| 14 | *SSU72* | 230085 | 230497 |
| 14 | *SLZ1* | 271876 | 272439 |
| 14 | *YNL195C* | 272926 | 273388 |
| 14 | *DUG3* | 282377 | 282932 |
| 14 | *MEP2* | 357876 | 358377 |
| 14 | *MLS1* | 406845 | 407223 |
| 14 | *DMA2* | 407461 | 407867 |
| 14 | *DBP2* | 416841 | 417224 |
| 14 | *INP52* | 425373 | 425687 |
| 14 | *LEU4* | 426182 | 426642 |
| 14 | *RPL9B* | 499869 | 500253 |
| 14 | *OCA2* | 518294 | 518670 |
| 14 | *COX5A* | 531846 | 532226 |
| 14 | *GPI15* | 558565 | 559030 |
| 14 | *NCE103* | 559917 | 560404 |
| 14 | *SIW14* | 575566 | 575958 |
| 14 | *HHF2* | 576757 | 577185 |
| 14 | *DOM34* | 630065 | 630470 |
| 14 | *DSE4* | 760402 | 760760 |
| 14 | *YNR068C* | 761348 | 761766 |
| 14 | *BSC5* | 762108 | 762605 |
| 15 | *YOL155W-A* | 29318 | 29736 |
| 15 | *HPF1* | 30970 | 31350 |
| 15 | *RPL25* | 82333 | 82714 |
| 15 | *TRM11* | 87587 | 88034 |
| 15 | *ZEO1* | 110298 | 110689 |
| 15 | *MHF1* | 159858 | 160315 |
| 15 | *GPD2* | 217936 | 218391 |
| 15 | *ARG1* | 219550 | 220066 |
| 15 | *AIM39* | 231342 | 231805 |
| 15 | *HTZ1* | 304904 | 305336 |
| 15 | *PLB3* | 306781 | 307200 |
| 15 | *GLO4* | 407737 | 408164 |
| 15 | *ETT1* | 426720 | 427234 |
| 15 | *RPL3* | 444942 | 445503 |
| 15 | *CYT1* | 447578 | 447960 |
| 15 | *KTR1* | 513340 | 513712 |
| 15 | *IDH2* | 580491 | 581054 |
| 15 | *MPC54* | 668123 | 668585 |
| 15 | *GAC1* | 669346 | 669728 |
| 15 | *DED1* | 669933 | 670494 |
| 15 | *HIS3* | 722155 | 722557 |
| 15 | *DED1* | 723762 | 724084 |
| 15 | *GEP3* | 724925 | 725377 |
| 15 | *ODC2* | 759101 | 759509 |
| 15 | *RPB8* | 761608 | 762000 |
| 15 | *DFR1* | 780246 | 780632 |
| 15 | *FSF1* | 832312 | 832728 |
| 15 | *MUM3* | 877230 | 877607 |
| 15 | *CPA1* | 882947 | 883368 |
| 15 | *MNE1* | 995469 | 995842 |
| 15 | *RAD17* | 1028176 | 1028593 |
| 15 | *RPS12* | 1040305 | 1040702 |
| 15 | *ALD4* | 1041882 | 1042271 |
| 15 | *FIT3* | 1059706 | 1060139 |
| 15 | *FIT2* | 1060220 | 1060705 |
| 16 | *DIM1* | 39564 | 39999 |
| 16 | *DIP5* | 41984 | 42383 |
| 16 | *FUM1* | 47979 | 48459 |
| 16 | *YAH1* | 74273 | 74657 |
| 16 | *CIN2* | 96718 | 97125 |
| 16 | *HSP82* | 97943 | 98332 |
| 16 | *USV1* | 109406 | 109913 |
| 16 | *PEP4* | 109930 | 110357 |
| 16 | *ODC1* | 111913 | 112125 |
| 16 | *SPO19* | 113009 | 113457 |
| 16 | *FAS2* | 113866 | 114254 |
| 16 | *AFT2* | 169874 | 170436 |
| 16 | *KIP2* | 260076 | 260523 |
| 16 | *ISU1* | 297785 | 298206 |
| 16 | *RPL5* | 303291 | 303693 |
| 16 | *CAR1* | 340238 | 340618 |
| 16 | *RPS9A* | 405034 | 405432 |
| 16 | *ERG10* | 498240 | 498798 |
| 16 | *AEP3* | 550996 | 551515 |
| 16 | *PDH1* | 558769 | 559176 |
| 16 | *GLN1* | 642506 | 643069 |
| 16 | *SPO24* | 645954 | 646375 |
| 16 | *BRR1* | 673098 | 673478 |
| 16 | *ROX1* | 680359 | 680748 |
| 16 | *TEF1* | 701143 | 701574 |
| 16 | *GRS2* | 701808 | 702217 |
| 16 | *CTR1* | 786276 | 786715 |
| 16 | *YLH47* | 786881 | 787302 |
| 16 | *NOC4* | 821669 | 822092 |
| 16 | *ASN1* | 823064 | 823437 |
| 16 | *NCE102* | 830114 | 830511 |
| 16 | *GPH1* | 862665 | 863031 |
| 16 | *AQY1* | 922108 | 922660 |

B. 3X-FLAG-Pch2-E399Q (Additional binding sites)

| **Chromosome** | **Gene** | **Start (bp)** | **End (bp)** |
| --- | --- | --- | --- |
| 1 | *TRN1* | 139446 | 139915 |
| 2 | *NUP170* | 80141 | 80628 |
| 2 | *HIS7* | 716867 | 717333 |
| 2 | *YBR284W* | 771275 | 771742 |
| 2 | *APE3* | 775680 | 776191 |
| 2 | *YBR296C-A* | 800505 | 801018 |
| 4 | *YDL183C* | 132189 | 132646 |
| 4 | *IDP1* | 335257 | 335831 |
| 4 | *NAT1* | 382634 | 383115 |
| 4 | *TRP1* | 462085 | 462516 |
| 4 | *ARO3* | 522362 | 522911 |
| 4 | *HSP42* | 806815 | 807416 |
| 4 | *SDH4* | 818119 | 818588 |
| 4 | *EBS1* | 866953 | 867438 |
| 4 | *RMD5* | 968452 | 969060 |
| 4 | *ATP17* | 1228628 | 1229154 |
| 4 | *YDR524C-B* | 1490585 | 1491064 |
| 5 | *RMD6* | 15627 | 16064 |
| 5 | *RIP1* | 107558 | 108129 |
| 5 | *GLC3* | 133411 | 133818 |
| 5 | *IRC22* | 151963 | 152437 |
| 5 | *RGI1* | 292205 | 292617 |
| 5 | *GCG1* | 504565 | 504981 |
| 5 | *BMH1* | 545930 | 546378 |
| 5 | *DMC1* | 548918 | 549326 |
| 6 | *EPL1* | 90495 | 90947 |
| 6 | *VTC2* | 133996 | 134552 |
| 6 | *DEG1* | 148322 | 148864 |
| 7 | *ATG1* | 162323 | 162826 |
| 7 | *YGL036W* | 428864 | 429308 |
| 7 | *HIP1* | 882900 | 883510 |
| 7 | *ADE3* | 907568 | 908137 |
| 7 | *MGA1* | 988941 | 989541 |
| 7 | *CWC22* | 1049348 | 1049883 |
| 7 | *YOR1* | 1058104 | 1058545 |
| 8 | *ECM29* | 40288 | 40754 |
| 8 | *QCR10* | 107938 | 108370 |
| 8 | *YAP1801* | 423049 | 423456 |
| 9 | *SUC2* | 38403 | 38852 |
| 9 | *NDC80* | 78857 | 79214 |
| 9 | *YIL060W* | 247858 | 248332 |
| 10 | *ERG20* | 106424 | 106873 |
| 10 | *MEF2* | 231751 | 232242 |
| 10 | *APL1* | 448287 | 448705 |
| 10 | *MHO1* | 453820 | 454244 |
| 10 | *CYC1* | 526395 | 526803 |
| 10 | *SFC1* | 609836 | 610299 |
| 10 | *RSF2* | 663746 | 664206 |
| 11 | *TRP3* | 38638 | 39191 |
| 11 | *NUP120* | 334400 | 334860 |
| 11 | *IXR1* | 382986 | 383458 |
| 11 | *PUT3* | 410388 | 410818 |
| 11 | *MET14* | 440122 | 440557 |
| 11 | *RSC4* | 454470 | 454922 |
| 11 | *KTR2* | 558918 | 559356 |
| 12 | *DNM1* | 150729 | 151320 |
| 12 | *RSC58* | 210338 | 210824 |
| 12 | *HAP1* | 646437 | 646937 |
| 12 | *GSY2* | 663397 | 663865 |
| 12 | *SKI2* | 919727 | 920071 |
| 12 | *ECM30* | 1011619 | 1012075 |
| 12 | *SST2* | 1042935 | 1043383 |
| 13 | *GTR1* | 28501 | 29108 |
| 13 | *TSL1* | 70861 | 71280 |
| 13 | *CAT2* | 192977 | 193526 |
| 13 | *MRPL39* | 252258 | 252736 |
| 13 | *MUB1* | 468476 | 469042 |
| 13 | *SPG4* | 483062 | 483492 |
| 13 | *GAT2* | 541904 | 542328 |
| 13 | *YMR206W* | 676281 | 676803 |
| 14 | *PCL1* | 89127 | 89533 |
| 14 | *WHI3* | 270459 | 270942 |
| 14 | *YNL190W* | 282438 | 282887 |
| 14 | *CUZ1* | 342776 | 343236 |
| 14 | *SMM1* | 655404 | 655877 |
| 14 | *ARE2* | 665461 | 666046 |
| 15 | *SHR5* | 110281 | 110691 |
| 15 | *PKH2* | 130900 | 131485 |
| 15 | *YSP3* | 331647 | 332256 |
| 15 | *HSP10* | 371228 | 371658 |
| 15 | *HUA2* | 849631 | 850118 |
| 15 | *YOR302W* | 882872 | 883474 |
| 15 | *EMT2* | 977650 | 978226 |
| 15 | *CIN1* | 993696 | 994153 |
| 16 | *TGS1* | 256233 | 256671 |
| 16 | *PXA1* | 274018 | 274601 |
| 16 | *PDR12* | 452150 | 452578 |
| 16 | *LEE1* | 456009 | 456442 |
| 16 | *ULP1* | 516449 | 516982 |
| 16 | *CIT3* | 556697 | 557099 |
| 16 | *ICL2* | 569331 | 569911 |
