## Supplementary Table 2 for "Active transcription and Orc1 drive chromatin association of the AAA+ ATPase Pch2 during meiotic G2/prophase"

| **Gene Ontology term** | **p-value** |
| --- | --- |
| Oxoacid metabolic process | 5.35e-16 |
| Carboxylic acid metabolic process | 5.54e-15 |
| Organic acid metabolic process | 1.47e-14 |
| Small molecule metabolic process | 6.01e-13 |
| Small molecule biosynthetic process | 1.77e-08 |
| Carboxylic acid biosynthetic process | 2.17e-08 |
| Single-organism metabolic process | 4.12e-08 |
| Organic acid biosynthetic process | 6.90e-08 |

List of GO terms associated with 3XFLAG-Pch2, generated by SGD GO term finder (molecular function) (<https://www.yeastgenome.org/goTermFinder)>, p-value was set to 0.01
