## Supplementary figures and images for "Active transcription and Orc1 drive chromatin association of the AAA+ ATPase Pch2 during meiotic G2/prophase"

### Supplementary Figure 1

#### Supplementary Figure 1

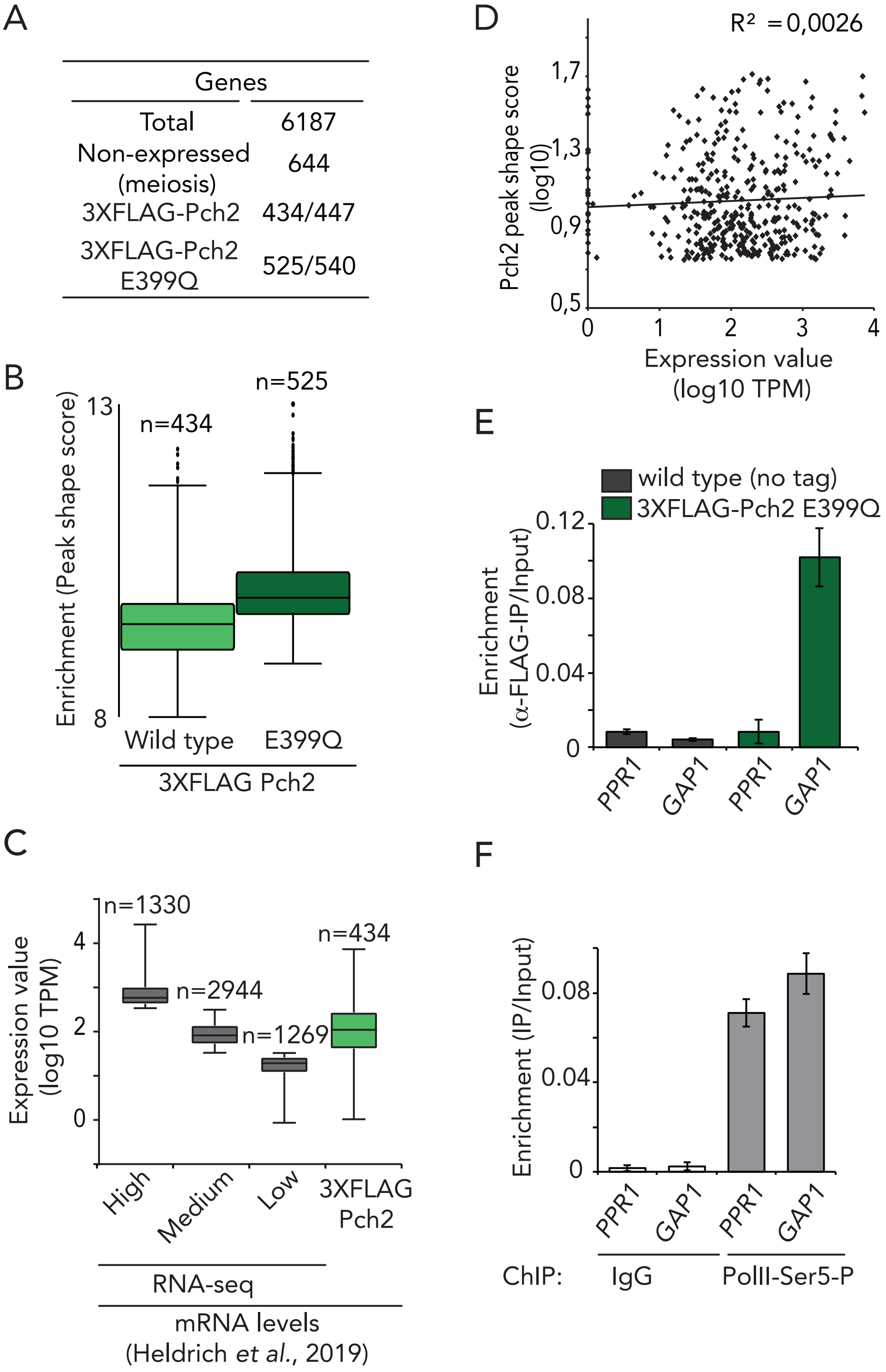

### Supplementary Figure 2

Supplementary Figure 2

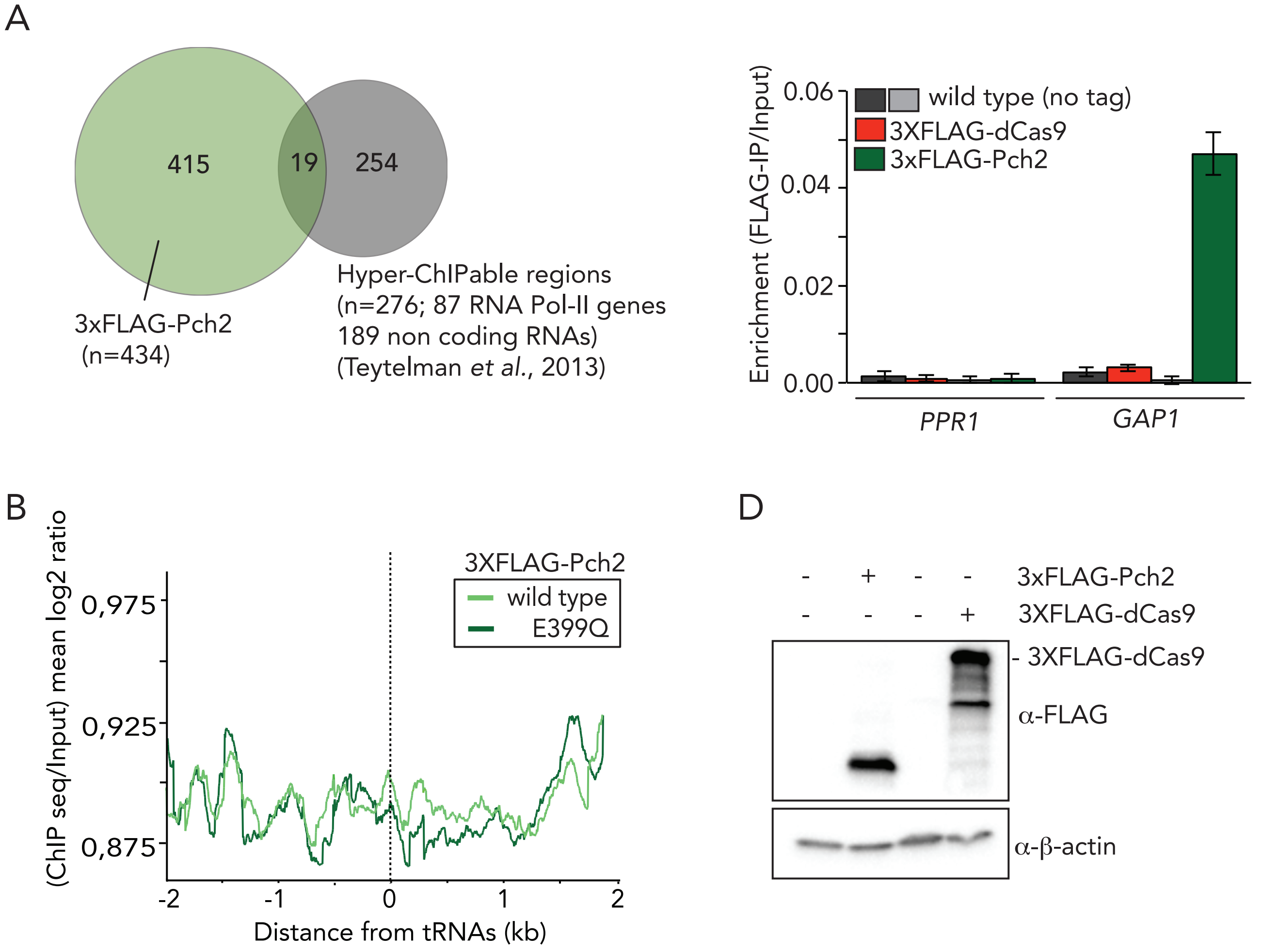

### Supplementary Figure 3

#### Supplementary Figure 3

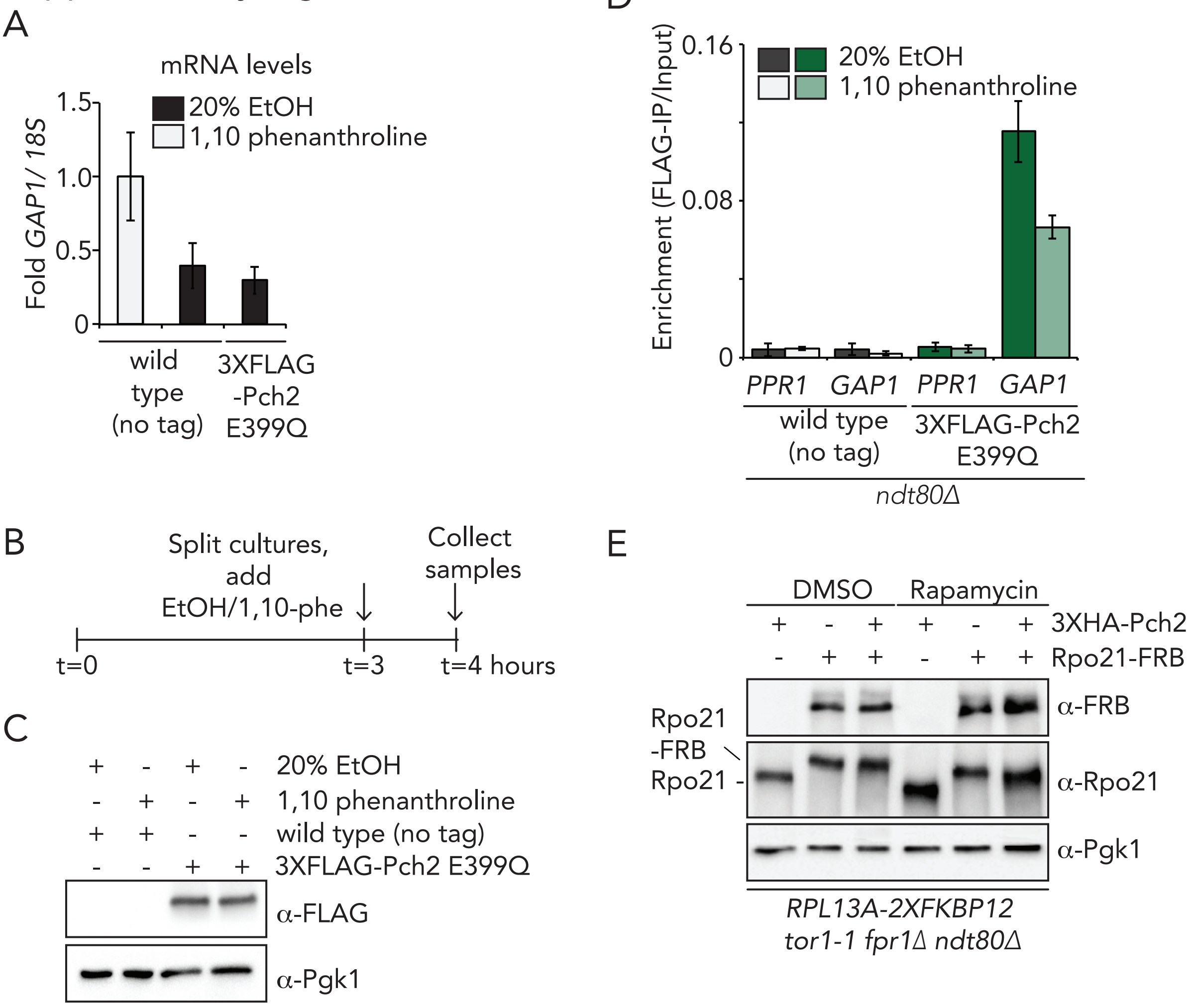

### Supplementary Figure 4

#### Supplementary Figure 4

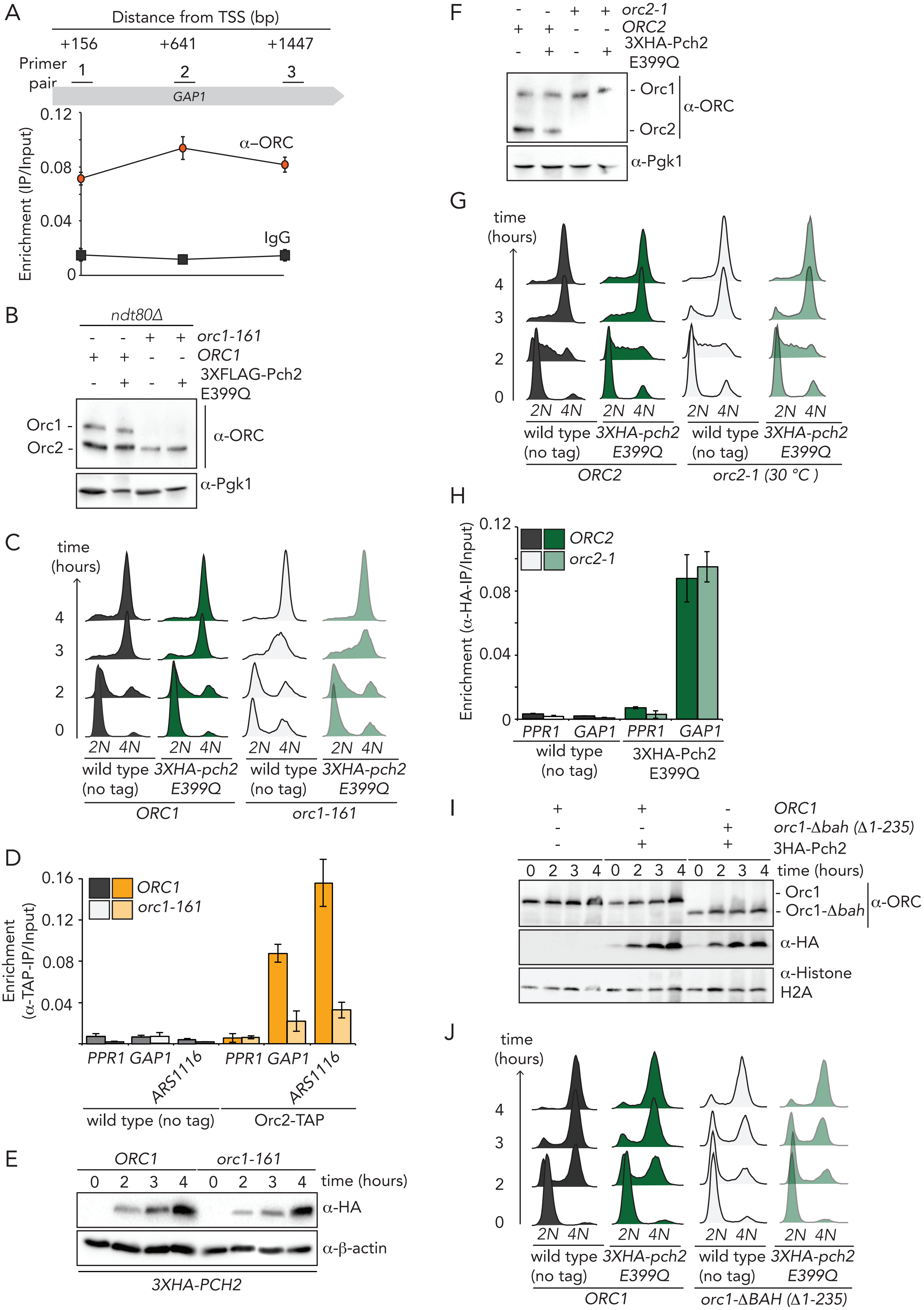

### Supplementary Figure 5

Supplementary Figure 5

A

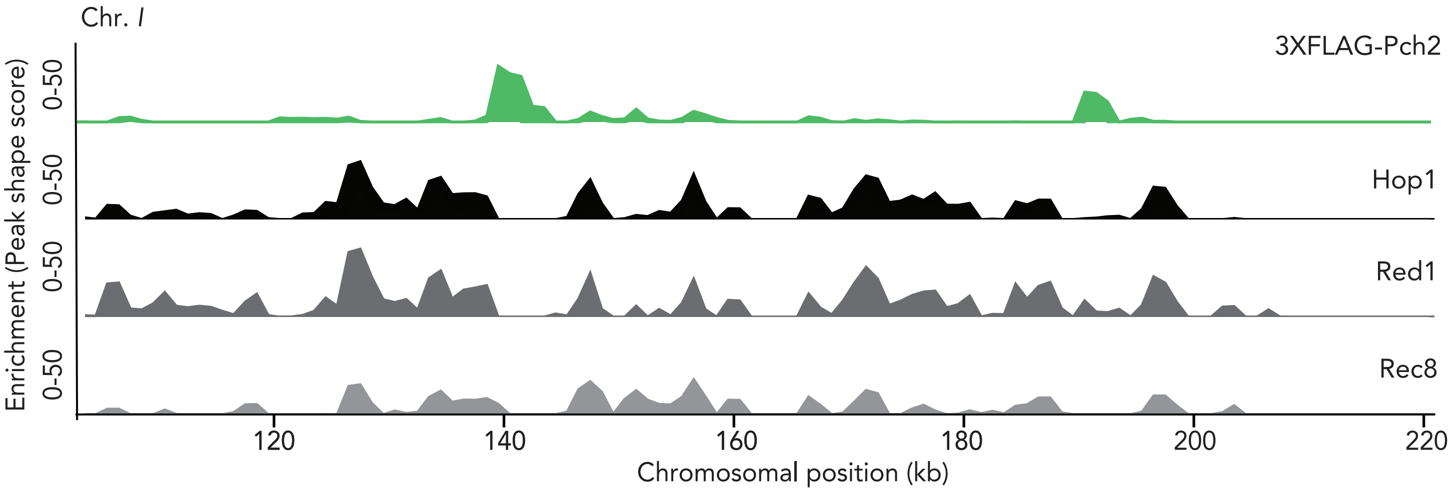


B

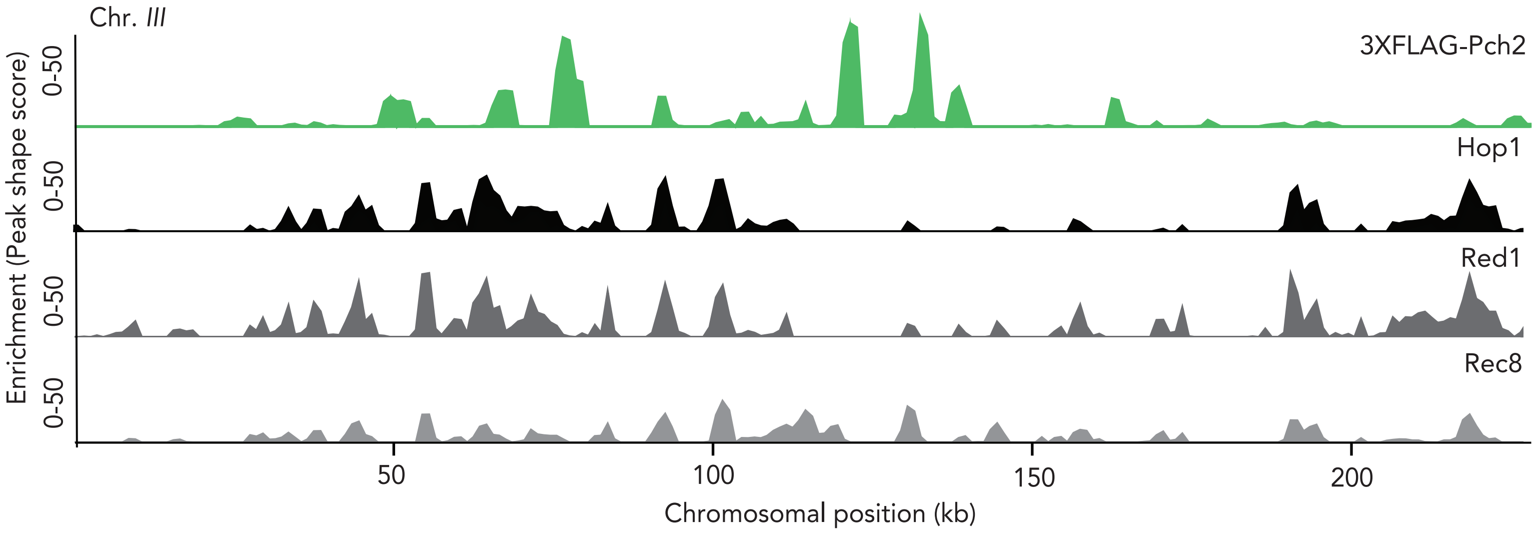

### Supplementary Figure 6

#### Supplementary Figure 6

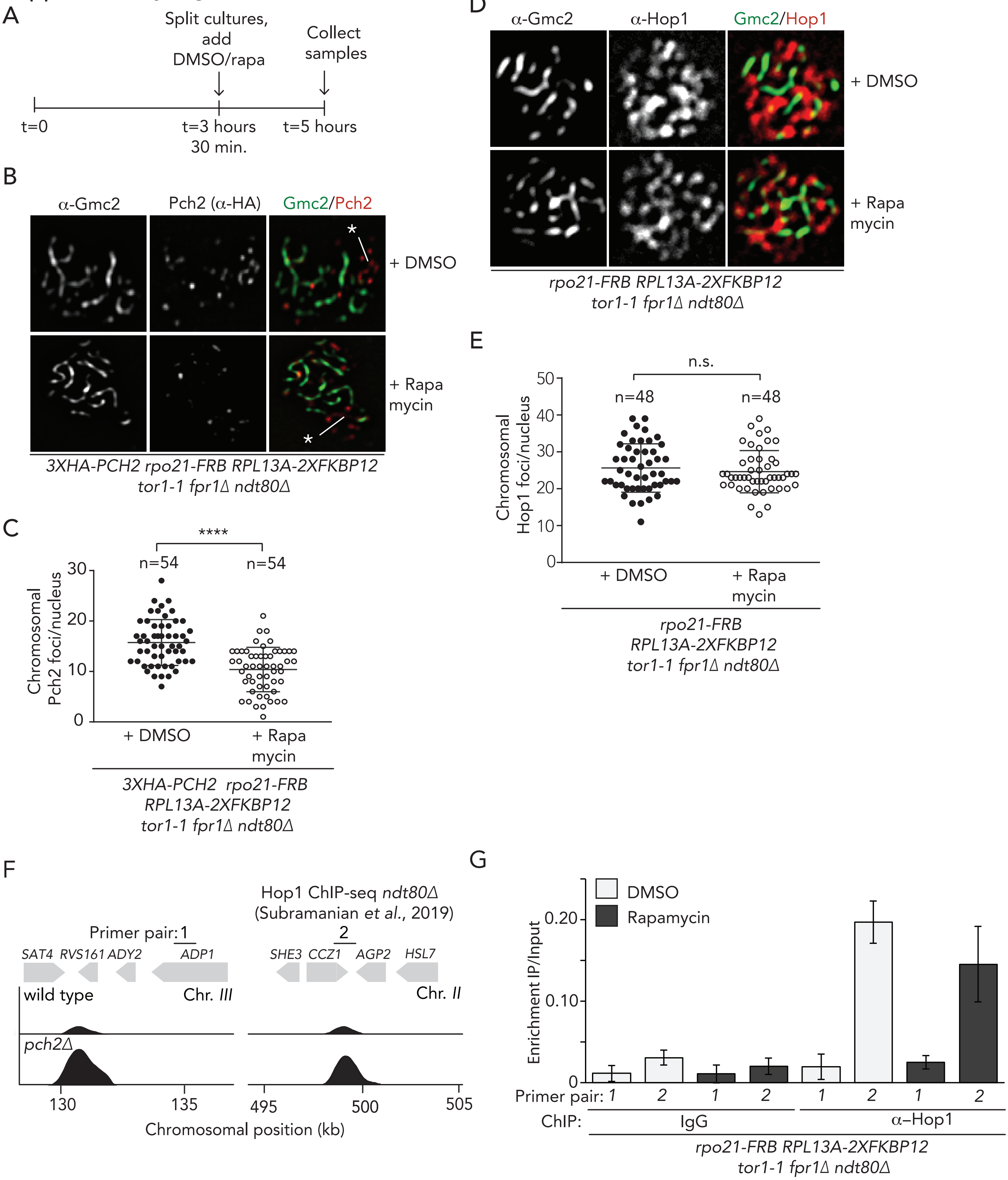

### Supplementary Figure 7

#### Supplementary Figure 7

A

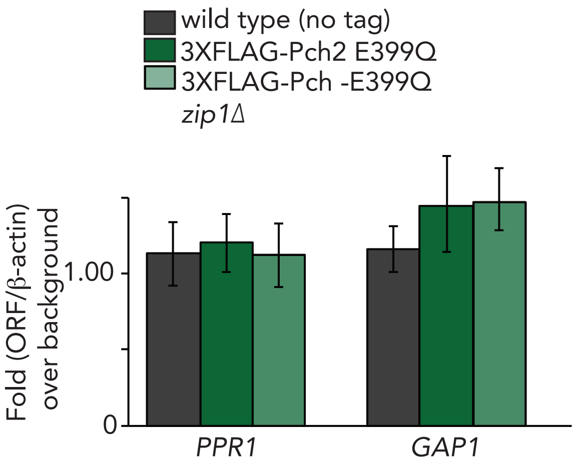


B

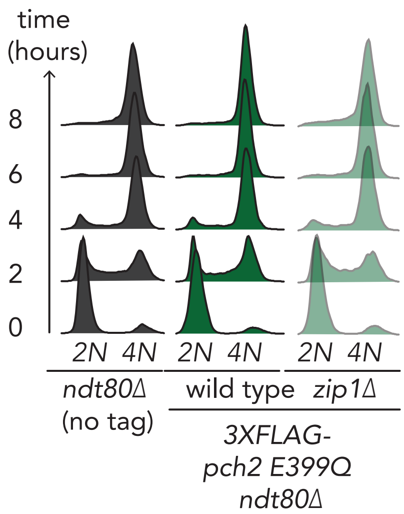


C

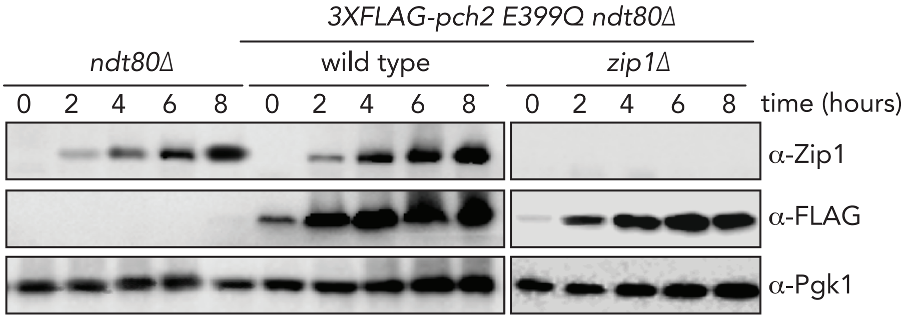
